# Deep computational analysis of human cancer and non-cancer tissues details dysregulation of eIF4F components and their interactions in human cancers

**DOI:** 10.1101/2020.10.12.336263

**Authors:** Su Wu, Gerhard Wagner

**Author notes:** Correspondence (G.W.).

## Abstract

Eukaryotic translation initiation complex (eIF4F) plays roles so diverse in human cancers as to complicate development of an overarching understanding of eIF4F’s functional and regulatory impacts across tumor types. Our analysis of large public data sets yielded several useful findings. *EIF4G1* frequently gains gene copies and is overexpressed to achieve characteristic stoichiometries with *EIF4E* and *EIF4A1* in cancers. Varied expressions among *EIF4F* components distinguish malignant from healthy tissues, regardless of tissue or cancer types. *EIF4G1* expression in particular correlates with poor prognosis. Tumors dysregulate biological “house-keeping” pathways typically regulated by cap-dependent initiation in healthy tissues, yet strengthen regulation of cancer-specific pathways in cap-independent contexts. In lung adenocarcinoma, altered interactions among eIF4F subunits are mechanistically linked to eIF4G1 phosphorylation. Tumors may select between cap-dependent and -independent mechanisms, through eIF4G1’s adaptable interactions with eIF4F subunits. Collectively, these results are an important advance towards a general model of translation initiation in cancer.

## INTRODUCTION

Protein synthesis supports cellular homeostasis and proliferation in healthy tissues, as well as growth and development of new organisms. Cancer development and progression also rely on protein synthesis, to support malignant behaviors such as tumor cell survival, angiogenesis, transformation, invasion, and metastasis. The essential step in protein synthesis is translation, the biochemical process by which ribosomes, as macromolecular machines, read mRNA nucleotide sequences, translate codons into accordant amino acids, and catalyze the formation of peptide bonds between amino acids to generate polypeptide chains. Translation activities are often stimulated by malignant cancer cells as part of an onco-sustenance strategy (Wu and Naar, 2019) that permits them to survive, proliferate, and metastasize under adverse (e.g. anaerobic and nutrient-deficient) conditions – conditions that generally suppress protein synthesis in normal cells (Taylor et al., 2009).

In eukaryotic contexts, the first step of translation, initiation, is a multi-step process in which an mRNA is activated, and a ribosome is recruited to that mRNA (Eisen et al., 1998; Shyamsundar et al., 2005). Altogether, translation initiation serves as a rate-limiting step for protein synthesis (Hershey et al., 2012; Jackson et al., 2010). The regulatory nexus for translation initiation is eukaryotic translation initiation factor 4F (eIF4F), a trimeric protein complex consisting of eIF4E, eIF4G, and eIF4A (Merrick, 2015). eIF4E binds to N^7^-methylated guanosine triphosphate (m^7^GTP) at the 5’ end of an mRNA strand (5’cap). eIF4G is a large scaffold protein that interacts with mRNA at the 5’ untranslated region (5’UTR) and multiple proteins including eIF4E, eIF4A, eIF4B, the eIF3 complex, poly(A)-binding protein (PABP), and mitogen-activated protein kinase-interacting kinases 1/2 (Mnk1/2) (Gross et al., 2003; Hinnebusch, 2014; Yanagiya et al., 2009). eIF4E and eIF4G anchor RNA helicase eIF4A to the 5’ UTR of mRNA, and stimulate eIF4A to unwind mRNA secondary structure to permit attachment of a 43S ribosome. This initiation mechanism is called “cap-dependent” because it requires both the presence and the binding of eIF4E and the 5’cap. Cap-dependent translation initiation (CDTI) is typical of most eukaryotic mRNAs. There are also cap-independent translation initiation (CITI) mechanisms, for which the 5’cap need not bind to eIF4E even if both are present, and indeed neither the 5’cap nor eIF4E is always strictly required (Pelletier and Sonenberg, 2019). In most cases (e.g. poliovirus, encephalomyocarditis, hepatitis A virus, and some cellular mRNAs), CITI relies on eIF4G and its binding to the 5’UTR at the internal ribosome entry site (IRES) to recruit ribosomes (Walsh and Mohr, 2011).

Prevailing thought holds that tumors frequently activate eIF4E to enhance CDTI in support of transformation, tumorigenesis and metastasis (Graff et al., 2008). In this model, the needs of cellular proliferation and metabolism require continuous stimulation of CDTI by environmental nutrients, growth factors and hormones, via the “mechanistic target of rapamycin” (mTOR) pathway (Saxton and Sabatini, 2017). When active, mTOR Complex 1 (mTORC1) phosphorylates the eIF4E Binding Protein (4E-BP), renders eIF4E free from 4E-BP inhibition, and thus promotes CDTI (Gingras et al., 1999; Gruner et al., 2016; Igreja et al., 2014; Sekiyama et al., 2015a). The postulated dependence of cancer on eIF4E or CDTI to sustain malignancy has guided the development of cancer therapies (Bhat et al., 2015). However, physiological stresses such as hypoxia, nutrient deprivation, and chemo and radiation treatments tend to deactivate mTORC1, and consequently to promote availability of hypo-phosphorylated 4E-BP to block eIF4E-eIF4G interaction and inhibit CDTI (Hara et al., 1998; Liu et al., 2006). In the context of such stresses, CDTI could only occur in the presence of sufficiently overabundant eIF4E. Consistently, our own prior work has demonstrated that hypoxia, as found in solid tumors, upregulates transcription of eIF4E by utilizing a hypoxia response element in eIF4E’s promoter region (Yi et al., 2013).

In malignant tumors, enablement of translation initiation under all (including physiologically stressful) circumstances is requisite for viability. Several cancer-specific mRNAs (e.g. vascular endothelial growth factor A (VEGFA), fibroblast growth factor 2 (FGF2), and hypoxia-inducible factor 1α (HIF1A)) reportedly have recourse to both CDTI and CITI (Badura et al., 2012). Mechanistic diversity of this sort threatens to confound development of therapies related to eIF4F (or eIF4E specifically). eIF4E is a dispensable subunit of the complex in CITI mechanisms, whereas eIF4G and eIF4A are necessary for both CDTI and (with rare exceptions) CITI. Mechanistic dissection of eIF4F in tumors, including a quantitative understanding of subunit stoichiometry and the influence of subunit stoichiometric variance upon malignant phenotypes as well as biological pathways, would clarify which translation initiation mechanisms are employed under physiological conditions. The prevalence and clinical relevance of each mechanism in cancers are not yet well understood but are critical to the mission of improving the success of cancer treatment.

Although direct observation of eIF4F protein activity would be the ideal investigative approach, modern proteomic data collection techniques (Wohlgemuth et al., 2015) like the “bottom-up” or “top-down” methods (Timp and Timp, 2020) do not allow direct, precise protein assignment and quantitation of eIF4F, particularly the 250-kDa large protein eIF4G1. There are no normalized quantitative protein data across cancer types from public datasets that are suitable for stoichiometry analyses of eIF4F. However, mRNA and associated proteins involved in translation are highly stable once synthesized; and in that special case mRNA abundance is a valid proxy for protein abundance (Schwanhausser et al., 2011). Positive correlations between RNA-Seq and proteomics data for eIF4F genes have been reported in breast invasive carcinoma (Mertins et al., 2016) and lung adenocarcinoma (Gillette et al., 2020). Quantitative RNA-Seq data for initiation factors from a large number of biopsies offer statistical power and clinical relevance that traditional wet-lab biochemistry studies seldom achieve (Hutter and Zenklusen, 2018).

We used quantitative RNA-Seq data from The Cancer Genome Atlas (TCGA) to infer the abundance and stoichiometry of eIF4F subunits, assess the influence of *EIF4F* gene expressions on malignancy and cancer progression, and determine correlation strengths of *EIF4F* genes; and thus to gain mechanistic insights into translation initiation in 33 common cancer types. In addition, we infer interaction among eIF4F subunits from phosphorylation status that promote those interactions using phosphor-proteomics data of lung adenocarcinoma from the Clinical Proteomic Tumor Analysis Consortium (CPTAC). Our analysis reveals dysregulation of eIF4F and eIF4E associated CDTI mechanism in tumors, and a possible role for eIF4G1 in tumor adoption of CITI when such dysregulation occurs.

## RESULTS

### Tumors frequently overexpress and gain gene copies of *EIF4G1*, but lose copies of *EIF4E* and *EIF4A1*

Genetic alterations such as point mutations and copy number variations (CNVs) are typical of cancers and often serve as important diagnostic tools (Loeb, 2011; Shlien and Malkin, 2009). Overexpression of *EIF4E, EIF4EBP1*, and *EIF4G1* has each been attributed to the amplification of gene copies in many cancer types including breast, head and neck, and prostate cancers (Jaiswal et al., 2018; Rutkovsky et al., 2019; Sorrells et al., 1998; Sorrells et al., 1999). Recurrent amplification suggests positive selection of a gene for consequential overexpression of its gene product in tumors. However, it is unclear whether all subunits are equally prone to gene amplification to achieve overexpression of the entire eIF4F complex. Therefore, we analyzed the CNVs of *EIF4E, EIF4A1, EIF4G1* and *EIF4EBP1* in 33 TCGA cancer types, and examined the association of CNVs among *EIF4F* subunits in all 10,845 TCGA tumors combined (**Figures 1A–1B** and **S1A–S1B**). We categorized CNVs into five statuses: amplification, gain (possible duplication), diploid, shallow deletion (possible heterozygous deletion), and deep deletion (possible homozygous deletion), according to the array-based DNA copy number data from TCGA (Mermel et al., 2011). We found that CNV statuses of *EIF4E, EIF4A1, EIF4G1* and *EIF4EBP1* are substantially dissimilar from each other across 33 cancer types. Duplication and amplification of *EIF4G1* occur in a high percentage of tumors from most cancer types, particularly in lung squamous cell carcinoma and ovarian serous cystadenocarcinoma (**Figure 1A**). Both duplication and deletion of *EIF4EBP1* are common in most tumors (**Figure S1A**). In contrast, heterozygous deletions of *EIF4E* or *EIF4A1* frequently occur in most cancer types, including lung squamous cell carcinoma and ovarian serous cystadenocarcinoma (**Figures 1B** and **S1B**). In all cancer types combined, the overall CNV statuses of *EIF4G1* and *EIF4E* differ among tumors (**Figure 1C**). We found common increase of *EIF4G1* gene copies: 7.74% of tumors contain amplification and 26.79% of tumors contain duplication. For *MYC*, even more starkly, 11.45% of tumors contain amplification and 38.32% contain duplication. We found similar frequencies of increase and decrease of *EIF4EBP1* gene copies: 21.03% of tumors contain duplication, 4.24% contain amplification, and another 23.42% contain heterozygous deletion. In contrast, we found the heterozygous deletion of *EIF4E* in 40% of tumors and *EIF4A1* in 30.21%, even more frequent than the heterozygous deletion of *PTEN* that occurs in 29.29%. We examined statistical associations between gene pairs for their CNV across all TCGA tumors (**Figure 1D**). We found a negative correlation coefficient (r = −0.21) for CNVs between *EIF4G1* and *EIF4E*, suggesting a weak association (i.e. a co-occurrence) between increase of *EIF4G1* and loss of *EIF4E*. Copy number losses of *EIF4A1* and *EIF4E* are weakly associated (r = 0.22).

**Fig. 1.**
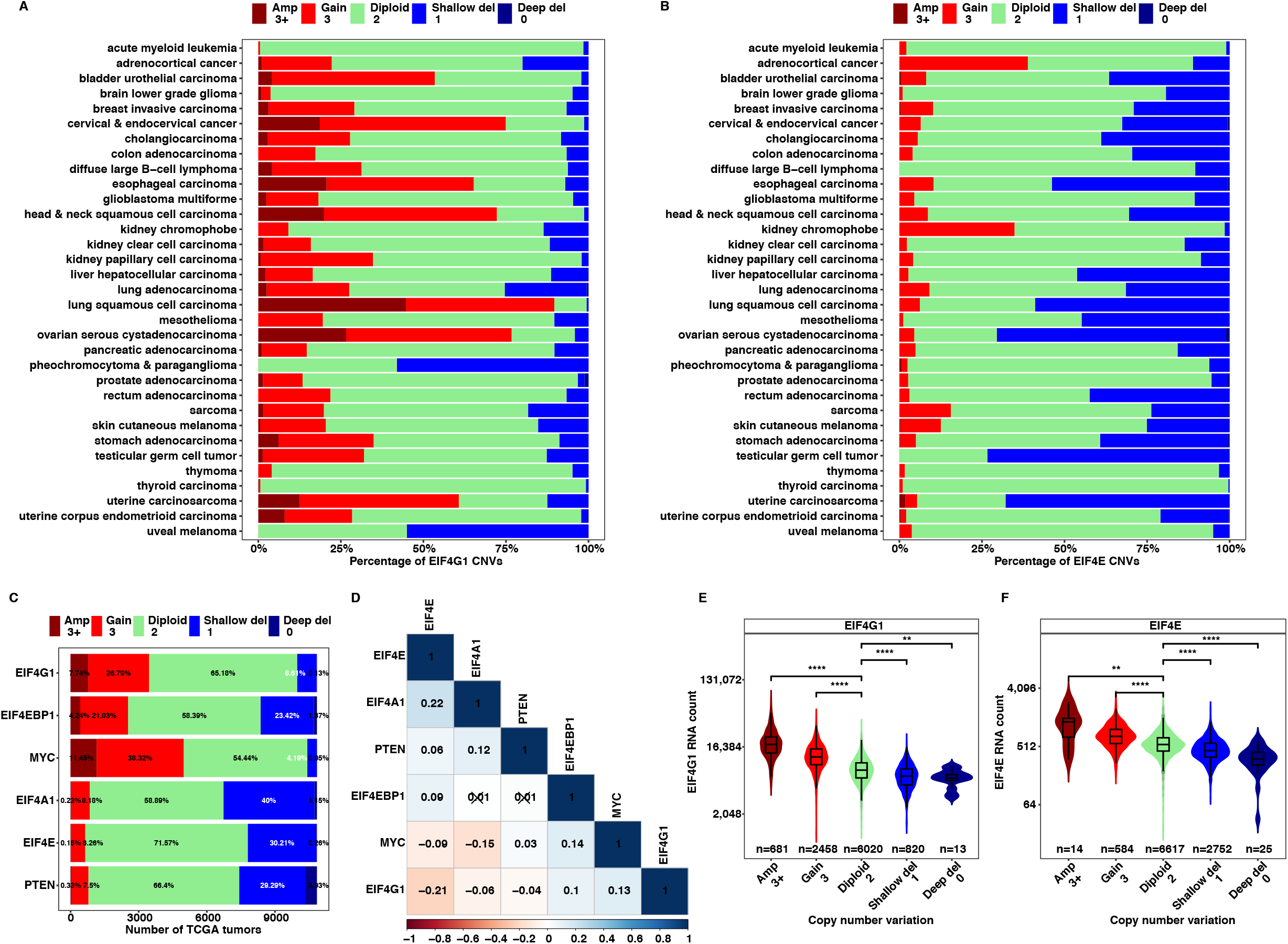
Tumors frequently overexpress and gain gene copies of *EIF4G1*, but lose copies for *EIF4E*. (**A** and **B**) The CNVs were estimated by GISTIC2 method using the whole genome microarray data. The estimated CNV values were categorized into five group with the following thresholds: 0, 1, 2, 3, 3+. These five groups represent deep deletion (possibly homozygous deletion), shallow deletion (possibly heterozygous deletion), diploid, low-level copy number gain (duplication), or high-level copy number gain (amplification) respectively. The stacked bar plots show the percentages of CNVs for *EIF4G1* and *EIF4E* in different TCGA tumor types. (**C**) The stacked bar plot shows the number of tumors with different CNVs for *EIF4G1, EIF4EBP1, MYC, EIF4E, EIF4A1* and *PTEN* in all TCGA tumors. The percentages of different CNVs were labeled on the bars. (**D**) The matrix plot shows the Pearson correlation coefficients between gene pairs for their CNV values across all TCGA tumors. p-value indicates the probability that the correlation may occur. The statistically insignificant correlation coefficients (p > 0.05) were crossed. (**E** and **F**) The associations between RNA expressions and CNV statuses were analyzed for *EIF4G1* and *EIF4E* in all TCGA tumor biopsies. The violin plots show the median values of RNA-Seq counts (transcripts per million) of *EIF4G1* and *EIF4EBP1* in the samples with different CNV statuses. The twotailed Student’s t tests were performed. ns, not significant; *P ≤ 0.05; **P ≤ 0.01; ***P ≤ 0.001; ****P ≤ 0.0001. Note: since CNVs were measured by Affymetrix array at the focal regions of genes, genes marked with deep deletion (deletion of two gene copies) may still express truncated mRNAs that can be detected by RNA-Seq.

To confirm that CNV status affects gene expression, we analyzed the relation between CNV status and mRNA expression for each *EIF4F* subunit. We found significant associations between RNA expression and CNV status for *EIF4G1* (**Figures 1E**), *EIF4E* (**Figures 1F**), *EIF4EBP1* (**Figures S1C**), *EIF4A1* (**Figures 1D**), *MYC* (**Figures S1E**), and *PTEN* (**Figures S1F**) across all 10,845 TCGA tumors. To verify the tumor-specific contribution of CNV status to gene expression, we used the average CNV from the adjacent normal tissues as the baseline control and analyzed the CNV ratio to that baseline for each tumor (**Figures S2A–S2D**). In most cancer types the average CNV ratios of *EIF4G1* and *MYC* are higher than 1, confirming the prevalence of gene duplication and amplification in tumors (**Figures S2A** and **S3A**). The average ratios for *EIF4EBP1* are close to 1 (**Figure S2C**), which is due to the similar prevalence of duplication and heterozygous deletion found in **Figure S1A**. The average CNV ratios for *EIF4E* (**Figure S2B**), *EIF4A1* (**Figure S2D**) and *PTEN* (**Figure S3B**) are lower than 1, suggesting predominant deletion. Finally, we analyzed mRNA expressions of *EIF4F* subunits in metastatic tumors, primary tumors, and adjacent normal biopsies from all TCGA cancer studies combined. *EIF4G1* expression is significantly elevated in metastatic and primary tumors vs. in adjacent normal biopsies (**Figure S3C**), likely due to the tumor-specific increase of gene copies (**Figures 1A** and **S2A**). Although significantly decreased in metastatic tumors, *EIF4E* expression is slightly increased in primary tumors in comparison to adjacent normal biopsies, (**Figure S3C**), despite the frequent heterogenous deletion of its gene in many cancer types (**Figures 1B** and **S2B**). However, *EIF4EBP1* and *EIF4A1* expressions are significantly elevated in metastatic and primary tumors vs. adjacent normal biopsies (**Figure S3C**), despite the similarly frequent duplication and deletion of *EIF4EBP1* gene (**Figures S1A** and **S2C**) and the prevalent loss of one *EIF4A1* gene copy (**Figures S1B** and **S2D**) in many cancer types. This puzzling result suggests some alternative regulatory influence over *EIF4E*, *EIF4EBP1* and *EIF4A1* expressions in cancers. In sum, *EIF4G1* overexpression is likely due to the amplification and duplication of gene copies in most tumors, suggesting *EIF4G1* is under positive selection.

### Tumors have unique stoichiometries of *EIF4G1:EIF4A1* and *EIF4G1*:(*EIF4E* + *EIF4EBP1*)

The CNV statuses alter mRNA expressions of eIF4F subunits in tumors, which likely perturb stoichiometric balance among subunits. Normalized protein quantitation data across cancer types are currently unavailable for stoichiometry analyses of eIF4F subunits, whereas RNA-Seq data quantify concentrations of mature mRNAs by normalizing sequencing counts of each gene to the total RNA counts and gene length. Thus, we used quantitative gene expression data from RNA-Seq to understand the abundance and stoichiometry of *EIF4F* mRNAs across pan-cancer types. We examined the relative abundances of *EIF4G1, EIF4A1, EIF4E*, and *EIF4EBP1*, in 33 cancer types from TCGA and 30 healthy tissue types from Genotype-Tissue Expression (GTEx). *EIF4G1* and *EIF4A1* are much more abundant than *EIF4E* and *EIF4EBP1* in all cancer types (**Figure 2A**) and all healthy tissue types (**Figure S4A**). Head & neck squamous cell carcinoma and lung squamous cell carcinoma have the highest average *EIF4G1* expression among all cancer types. However, upregulation of *EIF4G1* is not observed in normal lung samples (**Figure S4A**), which argues against any notion that *EIF4G1* overexpression in lung cancers originates from lung-tissue-specific gene expression. We reason that the *EIF4G1* overexpression in lung squamous cell carcinoma is linked to the cancer type and likely due to high incidence of its gene amplification and duplication (**Figure 1A**). The average expression of *EIF4E* does not vary widely across cancer types, with only two-fold difference between the highest and lowest ranking tumor types (**Figure 2A**). This result is consistent with the observation that most cancer types harbor the heterozygous deletion of *EIF4E* that can only affect up to 50% gene expression (**Figure 1B**).

**Fig. 2.**
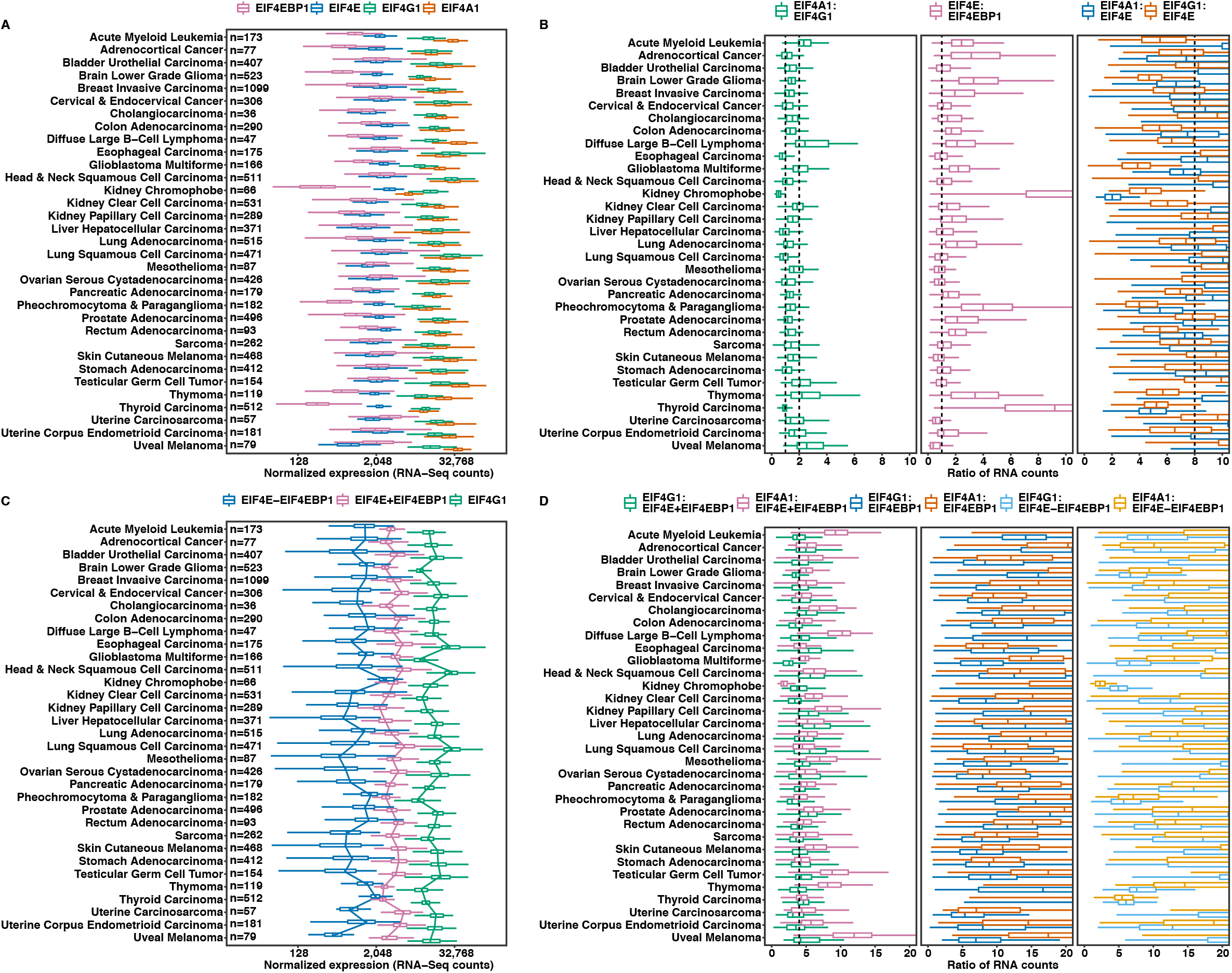
Tumors have unique stoichiometries of *EIF4G1:EIF4A1* and *EIF4G1*:(*EIF4E + EIF4EBP1*). (**A**) The expressions of *EIF4E, EIF4G1, EIF4A1* and *EIF4EBP1* was compared across different tumor types using the RNA-Seq data normalized for batch effects from 9,806 malignant tumors of 33 cancer groups in TCGA. The box and whisker plots represent gene expression (transcripts per million) from different cancer types in log_2_ scale. (**B**) The ratios of RNA expression were calculated in each sample of 9,806 malignant tumors and compared across 33 different tumor types as the box and whisker plot with a linear scale on the x-axis. The dash lines mark 1:1 and 2:1 ratio in the *EIF4A1:EIF4G1* panel, 1:1 ratio in the *EIF4EBP1:EIF4E* panel, and 8:1 ratio in the *EIF4G1:EIF4E* panel. (**C**) The sum expression of *EIF4E* and *EIF4EBP1* (*E+EBP1*), and the difference expression between *EIF4E* and *EIF4EBP1* (*E-EBP1*) were calculated in each tumor sample. They were compared to the *EIF4G1* expression across 33 TCGA tumor types. The trendlines connect the median values of boxplots. (**D**) The ratio between *EIF4G1* and *E+EBP1*, and the ratio between *EIF4G1* and *E-EBP1* were calculated within each tumor sample and compared across 33 cancer types. The dash line marks 4:1 ratio in the first panel.

We further calculated the ratios of *EIF4F* subunits within each sample, to infer possible stoichiometries in cancer or healthy tissue types. The average *EIF4A1:EIF4G1* ratios are between 1:1 and 2:1 in most tumor types (**Figure 2B**), but close to 1:1 in most healthy tissues except blood and muscle (**Figure S4B**). Blood cancers including acute myeloid leukemia and diffuse large B-cell lymphoma have the highest *EIF4A1*:*EIF4G1* ratios (> 2), which is likely due to the tissue-specific upregulation of *EIF4A1*:*EIF4G1* ratio (~ 4:1) in blood. Furthermore, across all cancer types combined, the average *EIF4A1:EIF4G1* ratio is 1:1 in adjacent normal tissues, but this ratio is particularly elevated in metastatic tumors (**Figure S3D**). These results reveal distinct stoichiometries of *EIF4A1* and *EIF4G1* in most healthy tissues (1:1) and in some tumors (2:1), a finding which hints preference of different compositions/topologies of the eIF4F complex by metastatic and by tumor-adjacent tissues.

The average *EIF4G1:EIF4E* ratios vary modestly across different healthy tissues (**Figures S4B** and **S5B**). An 8:1 ratio is observed in many tissue types with specific exceptions for brain, muscle, testis, liver and skin. This suggests a possible stoichiometry of *EIF4G1* and *EIF4E* (8:1). However, the average *EIF4G1:EIF4E* ratios vary greatly across different cancer types (**Figure 2B** and **S5A**), suggesting a lack of stoichiometry of *EIF4G1* and *EIF4E* in malignant tissues. In addition, we analyzed the *EIF4G1:EIF4E* ratios in all cancer types combined. *EIF4G1:EIF4E* ratios are significantly elevated in tumors vs. adjacent normal biopsies, with the highest ratios in metastatic tumors (**Figure S3D**). Given that *EIF4G1* overexpression is related to its CNVs in tumors, we reason that abolishment of the typical stoichiometry of *EIF4G1* and *EIF4E* is preferred in tumors.

The average *EIF4E* expression is comparable to *EIF4EBP1* in most cancer types (**Figure 2A**) but more abundant than *EIF4EBP1* in all healthy tissue types except pancreas (**Figure S4A**). The average *EIF4E1*:*EIF4EBP1* ratios are above 1 but vary greatly in most tumor types (**Figure 2B**) and healthy tissues (**Figure S4B**), indicating a lack of stoichiometry. However, across all combined cancer types the average *EIF4E1:EIF4EBP1* ratio is significantly higher in adjacent normal tissues than in tumors, (**Figure S3D**). These results suggest *EIF4E* is more abundant than *EIF4EBP1* in healthy tissues, but the difference between their expressions decreases dramatically in tumors. In metastatic tumors the average *EIF4E1*:*EIF4EBP1* ratio is below 1, suggesting *EIF4EBP1* is more abundant than *EIF4E*.

This altered ratio of *EIF4EBP1* and *EIF4E* may affect the availability of eIF4E in tumors, and thus perturb the stoichiometry *EIF4G1* and *EIF4E*. To examine this possibility, we calculated the difference between *EIF4E* and *EIF4EBP1* expressions (*E-EBP*) within each tumor (**Figure 2C**) or healthy tissue sample (**Figure S4C**), and we compared the expression of *E-EBP* to *EIF4G1*. The average ratios of *EIF4G1*:(*E-EBP*) and *EIF4A1*:(*E-EBP*) vary greatly across all cancer types (**Figure 2D**) and healthy tissues (**Figure S4D**), suggesting that the quantity of (*E-EBP*) does not help to define a useful stoichiometry. We further calculated the sum of *EIF4E* and *EIF4EBP1* expressions (*E+EBP*) within each tumor (**Figure 2C**) or healthy tissue sample (**Figure S4C**). The average expressions of *E+EBP* and *EIF4G1* show a parallel pattern across most cancer types (**Figure 2C**). The average *EIF4G1*:(*E+EBP*) ratios remain relatively constant around 4:1 in most cancer types (**Figures 2D** and **S5A**) but vary greatly across healthy tissues (**Figures S4D** and **S5B**). These results suggest a 4:1 stoichiometry of *EIF4G1* and *E+EBP* in cancers but a lack of stoichiometry in healthy tissues. Furthermore, although *EIF4G1* and *E+EBP* expressions are both significantly elevated in tumors vs. adjacent normal tissues of all combined cancer types (**Figure S3C**), the *EIF4G1*:(*E+EBP*) ratios remain 4:1 in the metastatic and primary tumors (**Figure S3D**). These results indicate that the 4:1 stoichiometry of *EIF4G1*:(*E+EBP*) is characteristic of tumors, regardless of their individual expression. One possible explanation is that the 4:1 stoichiometry promotes malignancy. Another is that tumors cease activities undertaken by healthy cells that drive stoichiometric variance. Either way, this tumor-characteristic stoichiometry hints at a tumor-characteristic molecular composition of the eIF4F complex.

### *EIF4G1* and *EIF4E* expressions are biomarkers that distinguish tumors from healthy tissues

CNVs triggered the alteration of mRNA expression levels and stoichiometries among *EIF4F* subunits in tumors, implying *EIF4F* genes may serve as biomarkers for tumor samples. However, since variability of single gene expression makes it less accurate to distinguish tumors from healthy tissues, we test whether collective expressions of *EIF4F* subunits distinguish tumors from healthy tissues. In addition to the four primary *EIF4F* genes, we also included *PABPC1, MNK1*, and *MNK2*, because the proteins coded by those genes biochemically interact with eIF4G1 and are deemed peripheral factors of the eIF4F complex. We applied principal component analysis (PCA) to the RNA-Seq data of those seven genes using the combined samples of primary and metastatic tumors from 33 cancer types and healthy samples from 30 tissue types (**Figures 3A** and **3B**). The first principal component (PC1) accounts for 38.1% of total gene expression variability, and PC2 accounts for 24.2% (**Figure 3A**); together they best account for the variability of total gene expression (62.3%). In **Figure 3A** overall, PC1 and PC2 are jointly capable of separating cancer cells from healthy ones, but neither one does so by itself. The nature of the PCs in **Figure 3A** is partly revealed by inspection of **Figures S6A–S6F**, in which the same PCs are preserved and the isolated subsets of samples are color-coded by type. PC1 strongly distinguishes between healthy brain tissues (magenta in **Figure S6D**), and metastatic skin cutaneous melanomas (brown in **Figure S6F**). PC2 strongly distinguishes between healthy brain tissues (magenta in **Figure S6D**) on the bottom, and healthy blood samples (yellow in **Figure S6D**). Most primary tumors from various cancer types cluster with significant overlap (**Figures S6B** and **S6E**). These results suggest that variability of *EIF4F* expression distinguishes tumors from healthy tissues, and also distinguishes healthy tissues from each other. We used correlation matrix plot to define gene expression variables for each PC. All seven genes contributed far more heavily to PC1 or PC2 than to any other PC; no gene contributed significantly to both PC1 and PC2 (**Figure 3B**). PC1 is composed primarily of *EIF4G1, EIF4A1, EIF4EBP1* and *PABPC1;* PC2 is composed of the *EIF4E, MKNK1* and *MKNK2*. We reason variability of all seven genes collectively contributes to distinguish tumors from healthy tissues. Joint *EIF4F* expressions serve as potential biomarkers for tumors or brain tissues.

**Fig. 3.**
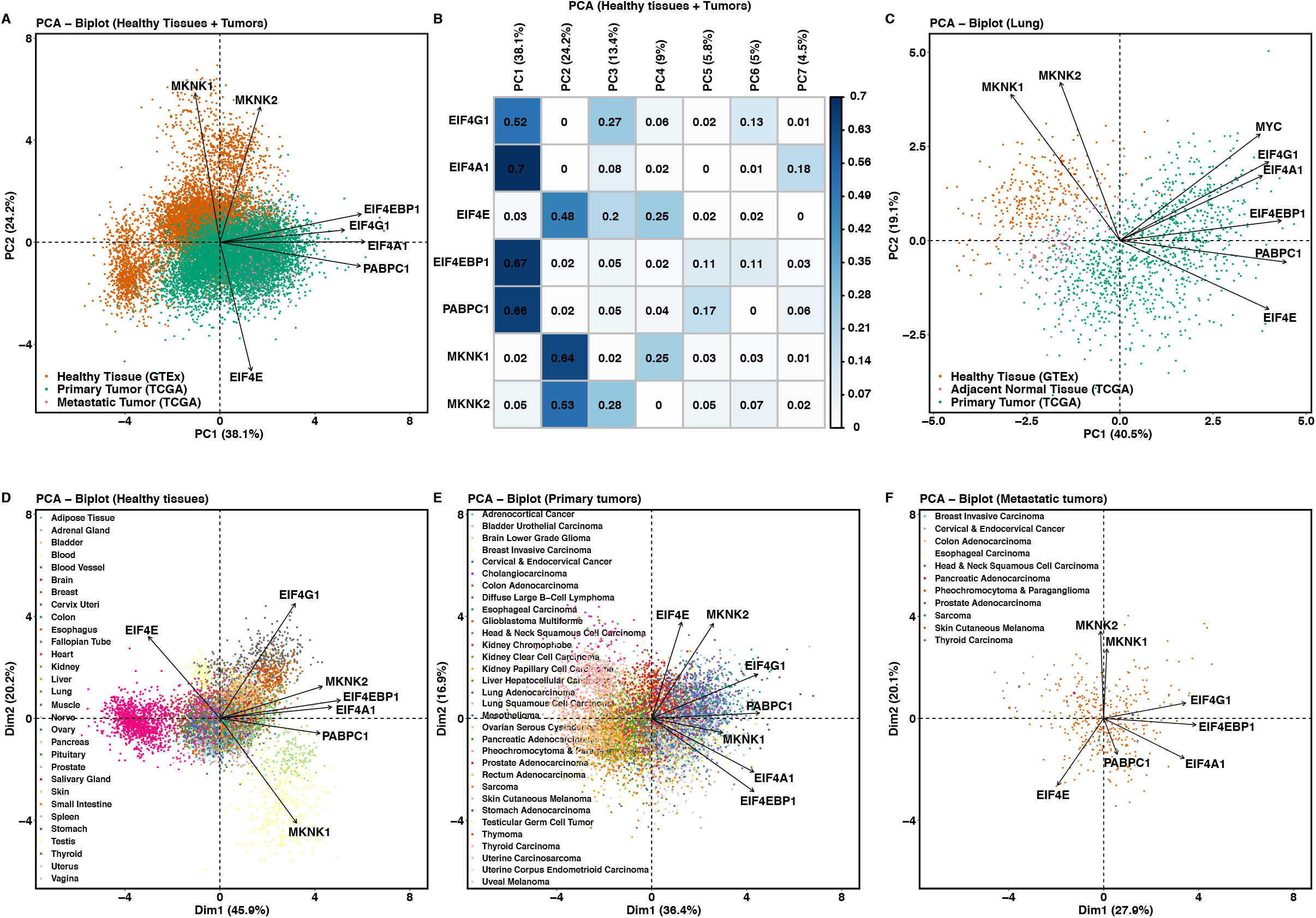
*EIF4G1* and *EIF4E* expressions distinguish tumors from healthy tissues. (**A**) PCA analysis using the standardized RNA expression of *EIF4G1*, *EIF4A1*, *EIF4E*, *EIF4EBP1*, *PABPC1, MKNK1* and *MKNK2* from 9,162 primary tumors and 392 metastatic tumors from TCGA and 7,388 normal tissue samples from GTEx. The gene expression levels were chosen as the variables in all PCAs and the sample types as the observations. The PCA biplot shows the first two principal components (PCs) along which the samples vary the most. Axis title shows the percentage of variances (squared loadings) explained by PC1 or PC2. Axis values show the PCA scores of individual samples. Arrows show the influence of each gene variable on the PCs. The sample types were not used to construct PCs and they were labeled in color afterwards on the PCA plot. (**B**) The correlation matrix plot for (**A**) shows the percent of variance in which each gene variable is explained by the PC – the values of squared cosines. Cosines refer to the correlation coefficients between the gene variables and PCs. The squared cosine reflects the representation quality of a gene variable on a PCA axis. (**C**) The PCA analysis using the standardized RNA expressions of *EIF4G1*, *EIF4A1*, *EIF4E*, *EIF4EBP1*, *PABPC1, MKNK1, MKNK2* and *MYC* from 1011 primary lung tumors (517 lung adenocarcinomas and 494 lung squamous cell carcinomas), 109 adjacent normal lung tissues from TCGA lung adenocarcinoma and lung squamous cell carcinoma study groups, and 287 healthy lung samples from GTEx. (**D**) The PCA analysis using the standardized RNA expressions of *EIF4G1*, *EIF4A1*, *EIF4E*, *EIF4EBP1*, *PABPC1, MKNK1, MKNK2* and *MYC* from 7,388 normal tissue samples of GTEx. The PCA biplot shows the first two PCs along which the samples vary the most. Arrows show the influence of each gene variable on the PCs. The tissue types were not used to construct PCs and individual samples were colored by their tissue types afterwards on the PCA plot. (**E**) The PCA analysis using the standardized RNA expressions of *EIF4G1, EIF4A1, EIF4E, EIF4EBP1, PABPC1, MKNK1, MKNK2* and *MYC* from 9,162 primary tumors of all TCGA cancer types. The PCA biplot shows the first two PCs along which the samples vary the most. Arrows show the influence of each gene variable on the PCs. The cancer types were not used to construct PCs and individual samples were colored by their cancer types afterwards on the PCA plot. (**F**) The PCA analysis using the standardized RNA expressions of *EIF4G1, EIF4A1, EIF4E, EIF4EBP1, PABPC1, MKNK1, MKNK2* and *MYC* from 392 metastatic tumors of all TCGA cancer types. The PCA biplot shows the first two PCs along which the samples vary the most. Arrows show the influence of each gene variable on the PCs. The cancer types were not used to construct PCs and individual samples were colored by their cancer types afterwards on the PCA plot. Note: 366 metastatic tumors are from TCGA skin cutaneous melanoma group.

To compare how variability of *EIF4F* expression differentiates sample types within tumor-only and healthy-only datasets, we performed three control PCAs on healthy tissue samples (**Figures 3D** and **S7A**), primary tumors of different cancer types (**Figures 3E** and **S7B**), and metastatic tumors (**Figures 3F** and **S7C**). We plotted the results 2-dimensionally using PC1 and PC2 (**Figures 3D–3F)**. Each biplot has its own individual PC1 and PC2 that define the sources of greatest variance within each individual data set. We found that, in healthy tissue samples, PC1 and PC2 account for 66.1% of all gene expression variability (**Figures 3D**). The healthy tissues separate widely along **Figure 3D**-PC1 and partially along **Figure 3D**-PC2, as multiple distinct blobs representing brain (cyan), pancreas (light green), muscle (grey), and blood samples (yellow). Because the expression variability of all seven genes is characteristic of **Figure 3D**-PC1, and *EIF4G1, EIF4E* and *MKNK1* are also characteristic of **Figure 3D**-PC2 (**Figure S7A**), we reason all *EIF4F* genes are useful in distinguishing different healthy tissue types. In contrast, the top two PCs account for only 53.3% of all expression variability in primary tumors (**Figure 3E**). Primary tumors from most cancer types form one big overlapping cluster at the center of PCA plot, yet a few cancer types such as head & neck squamous cell carcinoma (blue on the right side) and pheochromocytoma (pink on the left side) separate from each other horizontally along **Figure 3E**-PC1. **Figure 3E**-PC2 does not seem to play a significant role in separating categories of primary tumors. Because every gene except *EIF4E* has a significant contribution to **Figure 3E**-PC1 (**Figure S7B**), we reason that *EIF4E* is not as useful to distinguish cancer types as it is to distinguish healthy tissues. By implication, *EIF4E* plays a type-specific role in healthy tissues but not in tumors. Finally, we found that in metastatic tumors PC1 and PC2 account for merely 48% of all expression variability (**Figures 3F and S7C**). **Figure 3F**-PC1 and **Figure 3F**-PC2 fail to separate the samples in the PCA plot, which is expected because the metastatic samples are predominantly from one tumor type, skin cutaneous melanoma. These findings together indicate that *EIF4F* expression is well able to distinguish healthy tissue types from each other, and (particularly with regard to *EIF4E*) is much less able to do the same for tumors.

To understand the possible functional relevance of genes that contribute to the same PC, we repeated the PCA with additional control genes (**Figure S7D**): *MYC* – a gene relies on eIF4F for translation initiation (Nanbru et al., 1997), *JUN* – a gene relies on eIF3d but not eIF4F for translation initiation (Lee et al., 2016), and *YY1* – a gene transcriptionally regulates mRNAs with very short 5’UTR (Elfakess and Dikstein, 2008). The expression variability of *MYC*, *EIF4G1*, *EIF4A1, EIF4EBP1* and *PABPC1* primarily contributes to PC1, which conforms to the eIF4F usage for *MYC*’s translation initiation and *MYC* expression as a strong biomarker for malignancy. In contrast, *JUN* mainly contributes to PC4 by itself (**Figure S7E**), consistent with the eIF4F-independent mechanism for its translation initiation. Finally, *YY1* contributes less to PC1 and PC2 but more to PC3 (**Figure S7E**). Because mRNAs with very short 5’UTRs require little secondary structure unwinding and thus depend less on eIF4F complex, this observation also hints that most genes that correlate with *EIF4G1* or *EIF4E* contain long 5’UTRs.

Finally, we verified the finding that *EIF4F* gene expressions collectively serve as biomarkers for malignancy. We performed PCA on specific tumor types with their matched healthy tissues in order to assess the individual influence of each tissue type. The PCA plot shows wide separation between primary lung tumors (lung adenocarcinoma and lung squamous cell carcinoma) and healthy lung tissues along the PC1 and PC2 axes (**Figure 3C**). PCA of proteomics data from CPTAC shows similar separation between lung adenocarcinomas and paired normal lung tissues (**Figures S7F**). Thus, malignant and healthy samples are similarly well distinguished from each other by mRNA and protein expressions of *EIF4F* genes. We further performed PCA with the RNA-Seq data from the following tumor and tissue pairs: lower grade glioma and glioblastoma multiforme vs. healthy brain tissue (**Figure S8A**), breast invasive carcinoma vs. healthy mammary tissue (**Figure S8B**), colon adenocarcinoma vs. healthy colon tissue (**Figure S8C**), pancreatic adenocarcinoma vs. healthy pancreas tissue (**Figure S8D**), prostate adenocarcinoma vs. healthy prostate tissue (**Figure S8E**), skin cutaneous melanoma vs. healthy skin tissue (**Figure S8F**). Wide separations between malignant tumors and healthy samples are present in the PCA plots of all paired cases. Overall, these results confirm that *EIF4F* genes serve as biomarkers to distinguish tumors from healthy tissues in many cancer types.

### *EIF4G1* expression predicts poor prognosis in cancer patients

Given *EIF4F* genes are useful biomarkers to distinguish tumors from healthy tissues, the question arises as to whether all subunits are similarly useful predictors of cancer progression. Therefore, we first performed Kaplan-Meier (KM) analysis to compare the survival probabilities between the patients with high and low expression of each gene, considered independently from other genes. For each *EIF4F* gene, two patient groups were selected from 10,295 TCGA patients of all cancer types combined, based on the ranking of expression levels within their tumor biopsies. The survival probabilities of patients with high expressions (top 20%) of *EIF4G1* (**Figure 4A**), *EIF4A1* (**Figure 4B**), *EIF4EBP1* (**Figure 4D**), *MKNK1* (**Figure 4E**), *EIF4G2* (**Figure S9A**), *EIF4EBP2* (**Figure S9B**), *PABPC1* (**Figure S9C**), *MYC* (**Figure S9D**), *EIF4H* (**Figure S9F**) and *EIF3C* (**Figure S9G**) are significantly worse than those with low expression (bottom 20%), suggesting that high expression of each gene is disadvantageous for survival. In contrast, the survival probabilities of the patients with high expressions of *MKNK2* (**Figure 4F**) and *EIF4B* (**Figure S9E**) are significantly greater, suggesting that for each of those genes, high expression carries a survival advantage. We observed no significant difference of survival probabilities with regard to expression of *EIF4E* (**Figure 4C**), *EIF3E* (**Figure S9H**), *MTOR* (**Figure S9I**), *RPTOR* (**Figure S9J**), *RICTOR* (**Figure S9K**), or *RPS6KB1* (**Figure S9L**).

**Fig. 4.**
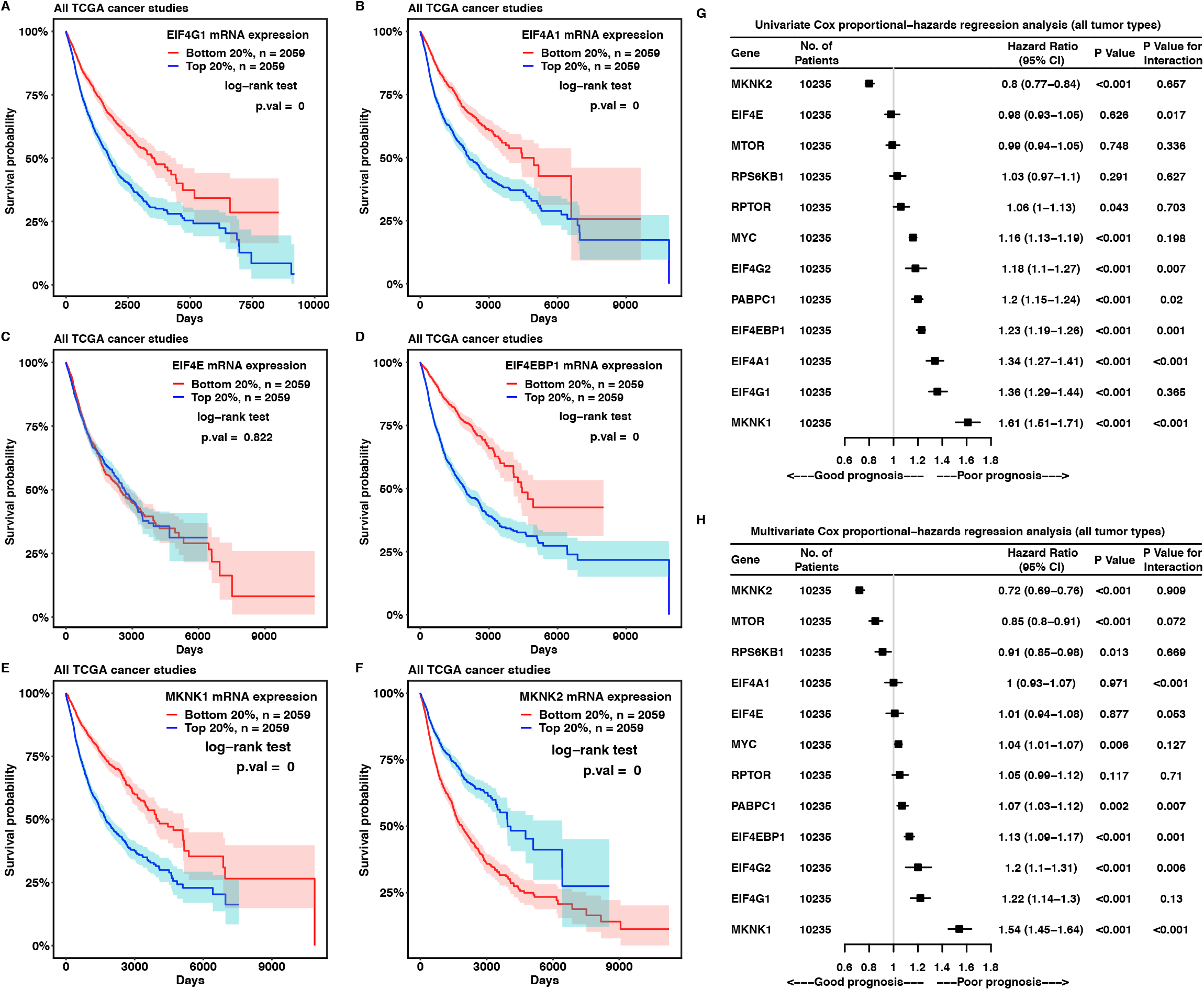
*EIF4G1* expression is an independent predictor for survival in cancer patients. (**A** to **F**) The Kaplan-Meier plots compare the survival probabilities of TCGA cancer patients according to the mRNA expressions of *EIF4G1* (**A**), *EIF4A1* (**B**), *EIF4E* (**C**), *EIF4EBP1* (**D**), *MKNK1* (**E**) and *MKNK2* (**F**) in their tumors. We ranked all 10,295 TCGA cancer patients based on the individual *EIF4F* gene expression from their tumor biopsies and selected two groups of patients with the top 20% or the bottom 20% of gene expression. The survival probability is the probability of an individual survives from the time origin (e.g. diagnosis of cancer) to a specified time. Differences in survival probabilities between two patient groups were assessed by the log-rank test. The shaded areas around the associated survival curves show the lower and upper bounds of the 95% confidence interval. (**G** and **H**) The univariate (**G**) and multivariate (**H**) Cox proportional-hazards regression models were examined for *EIF4F* expression in all 10,235 cancer patients from TCGA. P value indicates the statistical significance of association between gene expression and survival (i.e. a significant fit in Cox-PH model). Assumption of Cox proportional-hazards model was examined by Schoenfeld residues to test constant hazard ratio over time. A statistically significant P value for interaction indicates a violation on proportional-hazards model. CI, confidence interval.

To quantitatively evaluate how gene expression in tumors is related to patient survival, we then used the Cox proportional-hazards (PH) model to describe the relationships between hazard rate and expressions of one or more genes. The Cox PH regression model makes several assumptions, one of which is that hazard rates are proportional over time. We validated this proportionality by testing the interaction between gene expression and time. We first performed the univariate Cox-regression analyses that include a single gene expression as the dependent variable using the RNA-Seq data of all TCGA tumors combined (**Figure 4G**). We found that as the expression of *EIF4G1* (HR =□1.36, P value < 0.001) or *MYC* (HR =□1.16, P value < 0.001) increases, the chance of death increases, with no violation of the Cox model’s PH assumption (P values for interaction are 0.365 for *EIF4G1*, and 0.198 for *MYC*). These results indicate *EIF4G1* and *MYC* expressions each predict poor prognosis. *MKNK2* expression positively correlates with the survival length (HR =□0.8) with no violation of PH assumption (P value for interaction = 0.657) (**Figure 4G**), suggesting *MKNK2* expression predicts good prognosis. Although expressions of *MKNK1, EIF4A1, EIF4EBP1, PARPC1* and *EIF4G2* are each positively associated with survival length in the regression model, their PH assumptions are significantly violated and thus whether their gene expressions in tumor are related to patient survival remains inconclusive. Finally, to compare the impacts of multiple gene expressions simultaneously on patient survival, we performed the multivariate Cox regression to model death rate and expressions of twelve genes relevant to translation initiation (**Figure 4H**). Also here, *EIF4G1* expression significantly correlates with poor prognosis (HR =□1.22) with no violation of PH assumption (P values for interaction = 0.13), suggesting *EIF4G1* expression best predicts poor prognosis among all twelve genes. In contrast, the expressions of *MKNK2* (HR =0.72) and *MTOR* (HR =0.85) significantly correlate with good prognosis, with no violation of PH assumptions. In sum, *EIF4G1* expression significantly correlates with poor prognosis in the KM, univariate Cox and multivariate Cox analyses, indicating its critical role in disease progression of all analyzed cancer types.

To validate our findings in an individual cancer type, we performed KM analysis on 517 lung adenocarcinoma patients. We found the survival probabilities of the patients with high expressions of *EIF4G1* (**Figure S10A**), *EIF4A1* (**Figure S10B**), and also *EIF4E* (**Figure S10C**) are significantly worse than those with low expression, suggesting that high expression of each gene is disadvantageous for survival. However, we observed no significant difference of survival probabilities in patient groups regarding *EIF4EBP1* (**Figure S10D**), *MKNK1* (**Figure S10E**), and *MKNK2* **(Figure S10F**) expression. Univariate Cox-regression analyses indicate that the expressions of *EIF4G1* (HR = □1.54), *EIF4A1* (HR =□1.4) and *PARPC1* (HR =□1.3) are each negatively associated with survival length, with no violations of PH assumptions in the regression model (**Figure S10G**). Multivariate analysis on the twelve genes collectively (**Figure S10H**) indicate that only *EIF4G1* expression is negatively associated with survival length (HR =□1.78) with no violation of PH assumption. Thus, among the three analyses (KM, univariate Cox, and multivariate Cox), multiple translation initiation factors exhibit a degree of correlation with poor prognosis in lung adenocarcinoma; but *EIF4G1* alone exhibits correlation in all three.

### Tumors dysregulate biological pathways that correlate with *EIF4G1* and *EIF4E* expressions in healthy tissues

The notion that translation initiation subunits have different effects on cancer progression demands exploration of an underlying biological basis. Since genes that exhibit correlating expression usually serve biological processes of overlapping functions, we performed correlation analysis to identify genes dependent upon each *EIF4F* subunit. For each *EIF4F* gene (*EIF4E, EIF4G1, EIF4A1*, or *EIF4EBP1*), we correlated its expression with all cellular gene transcripts (58,582 in total) from either TCGA or GTEx RNA-Seq data. We calculated 58,582 Pearson’s correlation coefficients between each *EIF4F* gene and all gene transcripts across 10,323 tumor samples, and we focused on genes with positive correlations (posCORs, r > 0.3) or negative correlations (negCORs, r < −0.3). Similarly, we identified posCORs and negCORs for each *EIF4F* gene across 7,414 healthy tissues. In general, healthy tissues contain more positively correlating genes (posCORs) of *EIF4E, EIF4G1, EIF4A1* or *EIF4EBP1* than tumors (**Figure S11A**). Healthy tissues contain the highest number of posCORs (12,765) for *EIF4E*. Tumors have far fewer posCORs for any gene, the highest of which (2,601) is for *EIF4G1*. Healthy tissues and tumors alike contain fewer negatively correlating genes (negCORs) than posCORs (**Figures S11B–S11D**). In healthy tissues, *EIF4A1* and *EIF4G1* share most posCORs (3,170 + 2,814 + 1,118 + 272 = 7,374) (**Figure 5A**); *EIF4E* shares more than half of its posCORs with both *EIF4G1* and *EIF4A1;* all four genes share 2,814 posCORs. In tumors, *EIF4E, EIF4G1* and *EIF4A1* share only 50 posCORs; four genes share zero posCORs (**Figure 5B**). These results suggest that healthy tissues – and not tumors – have largely overlapping posCORs for *EIF4F* genes. The posCORs and negCORs that are common for *EIF4F* genes from healthy tissues or from tumors are shown in a clustered heat map, where their correlation strengths are compared (**Figure 5C**). We used hierarchical clustering to group samples based on the similarity of their correlating genes. We found that correlating genes from the same sample types tend to be clustered together. The posCORs for *EIF4F* genes from healthy tissues have much higher correlation coefficients than those from tumors (**Figure 5C**), implying stronger regulatory functions of *EIF4F* over those posCORs in healthy tissues.

**Fig. 5.**
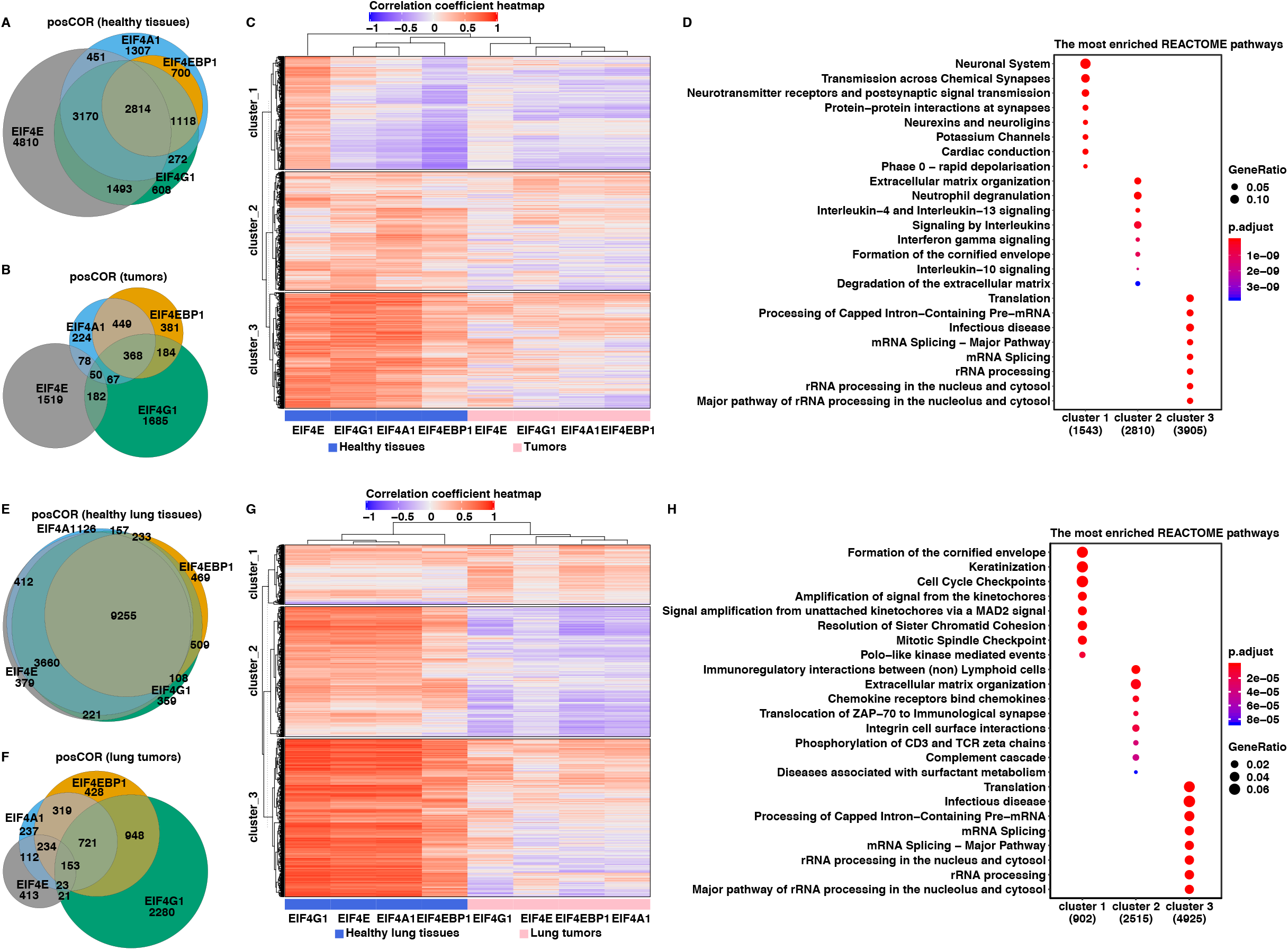
Tumors dysregulate biological pathways that correlate with *EIF4G1* and *EIF4E* expressions in healthy tissues. (**A** and **B**) The Pearson’s correlation coefficients between *EIF4F (EIF4E, EIF4G1, EIF4A1*, or *EIF4EBP1*) and other 58,582 genes were calculated separately in 10,323 TCGA tumor samples from different cancer types, or in 7,414 GTEx healthy samples from different tissue types, using the Toil Recomputed RNA-Seq datasets. The genes that have significant positive correlations (posCORs, r > 0.3) or negative correlations (negCORs, r < −0.3) were selected for analysis. The Venn diagrams show the overlapping posCORs among *EIF4F* in the healthy tissue samples (**A**) or in the tumors (**B**). (**C**) The heatmap shows the correlation strengths of posCORs and negCORs for *EIF4E, EIF4G1, EIF4A1* and *EIF4EBP1* in healthy tissues and in tumors. Each column represents an *EIF4F* gene from its belonged sample types: tumors or healthy tissues. Each row represents a correlating gene to the corresponding *EIF4F* gene. The color and intensity of each heatmap box represents the value of Pearson’s correlation coefficient for a correlating gene. Red represents posCORs and blue represents negCORs. White represents genes with no correlation with *EIF4F*. The dendrogram on the top shows the hierarchical relationship between the columns. The height of the dendrogram indicates the order in which the clusters were joined. The rows were ordered by K-means clustering to partition the dataset into three non-overlapping subgroups. (**D**) The dot plots show the enriched pathways in three clusters (K-means) from the heatmap (**C**) by REACTOME pathway analysis. The top 8 (most significantly) enriched pathways of each cluster were plotted. (**E** to **H**) posCORs and negCORs of *EIF4F* were separately identified from 1,122 lung tumors of lung squamous cell carcinoma (LUSC) and lung adenocarcinoma (LUAD) in TCGA, or from 287 healthy lung tissues in GTEx. The genes that have significant positive correlations (posCORs, r > 0.3) or negative correlations (negCORs, r < −0.3) were selected for analysis. The Venn diagrams show the overlapping posCORs among *EIF4F* from healthy lungs (**E**) or lung tumors (**F**). (**G**) The heatmap shows the correlation strengths of posCORs and negCORs for *EIF4E, EIF4G1, EIF4A1* and *EIF4EBP1* in healthy lungs and in lung tumors. (**H**) The dot plots show the enriched pathways in three clusters (K-means) of the heatmap (**E**) by REACTOME pathway analysis.

We used partitioning to classify correlating genes into three groups based on the data similarity (**Figure 5C**) and performed pathway enrichment analysis on the correlating genes from each cluster (**Figure 5D**). The genes grouped in “cluster one” have positive correlations with *EIF4E* in healthy tissues but not in tumors. Their biological functions are involved in the neuronal function-related pathways. The “cluster two” genes have moderate positive correlations with *EIF4E*, *EIF4G1* and *EIF4A1* in healthy tissues, but their correlation strengths became much weaker – particularly for *EIF4E* – in tumors. Their biological functions are involved in extracellular matrix organization and signaling pathways by interleukins. The “cluster three” genes have strong positive correlations with *EIF4E*, *EIF4G1* and *EIF4A1* in healthy tissues, but – as with cluster two – their correlation strengths become much weaker in tumors. Their biological functions are involved in house-keeping pathways including translation, pre-mRNA processing (splicing), and ribosomal RNA (rRNA) processing, which are shown to connect to translation initiation by previous studies (Lejeune et al., 2002). In sum, the dramatically decreased correlation strengths of posCORs in tumors suggest dysregulation of those posCORs and of related biological pathways by *EIF4F* genes. Each *EIF4F* gene has dramatically different correlation strength on the same posCORs in tumors, which we regard as the likely basis for the varying prognostic effects among *EIF4F* gene expressions on cancer patients (**Figures 4A** and **4C**).

To validate our finding in a specific cancer type with matched healthy tissue, we performed correlation analyses on *EIF4F* in lung tumors (lung squamous cell carcinoma and lung adenocarcinoma), and in healthy lung tissues. Normal lung tissues contain more positive correlation genes (posCORs) of *EIF4E*, *EIF4G1, EIF4A1* or *EIF4EBP1* than lung tumors (**Figure S11E**). There are fewer negCORs than posCORs for *EIF4E*, *EIF4G1*, *EIF4A1*, and *EIF4EBP1* in healthy lung tissues and in lung tumors (**Figures S11F–S11H**). In healthy lung tissues, *EIF4E, EIF4A1* and *EIF4G1* shared most of their posCORs (**Figure 5E**) and a total of 9,255 posCORs were shared by all four genes. These data suggest *EIF4E*, *EIF4G1* and *EIF4A1* have mostly overlapping functions in healthy lungs. However, in lung tumors, *EIF4E*, *EIF4G1* and *EIF4A1* share only 176 (153 + 23) posCORs; and all four genes share a mere 153 posCORs (**Figure 5F**). These results suggest lung tumors have limited overlapping posCORs among *EIF4F* genes. We used a clustered heat map to visualize posCORs and negCORs that are commonly regulated by *EIF4F* genes from healthy lungs or from lung tumors. The correlating genes from the same sample types tend to be clustered together. The same posCORs have much higher correlation coefficients in healthy lungs than in lung tumors (**Figure 5G**), implying stronger regulatory functions of *EIF4F* in healthy lungs. These results suggest that our interpretation of the full dataset is also suitable for a specific match of cancer and healthy tissue: *EIF4F* genes seem to have specific regulatory roles in healthy lungs that are greatly diminished in lung tumors.

We used partitioning to classify correlating genes in three clusters (**Figure 5G**) and performed pathway enrichment analysis (**Figure 5H**). The “cluster one” genes have weak positive correlations with *EIF4E*, *EIF4G1, EIF4A1*, and *EIF4EBP1* in healthy lungs. In contrast, those genes have modest positive correlations with *EIF4G1, EIF4A1* and *EIF4EBP1*, but no correlation with *EIF4E* in lung tumors. Their biological functions are involved in keratinization (formation of cornified envelope) – a pathological process associated with tumor progression in lung squamous cell carcinoma (Park et al., 2017) – and cell cycle regulation. The strengthened correlations of “cluster one” genes in tumors suggest that *EIF4F* genes reinforce their regulation of biological pathways that benefit malignancy. The “cluster two” genes have moderate positive correlations with all four genes in the healthy lungs, but have negative correlations with *EIF4G1, EIF4A1* and *EIF4EBP1* in lung tumors. Their biological functions were involved in extracellular matrix organization. The “cluster three” genes have strong positive correlations with all four genes in healthy lungs, but weak positive correlations in lung tumors. Their biological functions are involved in the house-keeping pathways such as translation and mRNA processing functions. A reasonable interpretation is that lung tumors generally diminish the traditional regulatory role of *EIF4F* genes over the house-keeping pathways, and repurpose *EIF4G1* and *EIF4A1* to regulate certain pathological pathways. The finding that *EIF4G1* and *EIF4E* vary in their regulatory strengths on pathological pathways is in line with their different prognostic effects on lung cancer patients (**Figures S10A** and **S10C**).

### Phosphorylation of a regulatory region in eIF4G1 is elevated in lung adenocarcinoma

Phosphorylation of eIF4F subunits reportedly regulates subunit interactions and CDTI. To understand the molecular basis of varying regulatory strengths of eIF4F subunits in tumors, we compared the location and relative abundance of eIF4E, eIF4A1, eIF4G1 and 4-EBP1 phosphorylation in tumors vs healthy tissues. We analyzed the relative abundances of peptides and phospho-peptides of eIF4F subunits, in 109 lung adenocarcinomas of four pathological stages and 102 matched normal adjacent tissue samples from CPTAC. The average abundance of eIF4G1 protein is significantly higher in lung adenocarcinomas from stages I to IV, than in normal lung tissues (**Figure 6A**). The phosphorylation of Serine and Threonine is found at many eIF4G1 sites, including Thr212, Thr214, Thr218, Ser503, Thr654, Ser711, Ser1035, Ser1068, Thr1080, Ser1084, Ser1087, Ser1099, Ser1151, Ser1152, Ser1201, Ser1216, Thr1218, Ser1238 and Ser1245 (**Figures 6B–6N** and **S12A–S12M**). **Figures 6B–6N** present the noteworthy phospho-peptides that are identified in more than 10% of all samples. The average abundance of eIF4G1 protein in lung adenocarcinomas (regardless of cancer stage) is 1.41 times that in normal adjacent tissues, whereas the average abundance of most phospho-peptides in lung adenocarcinomas (regardless of cancer stage) is more than twice that in normal adjacent lung tissues. These results demonstrate that lung tumors have higher eIF4G1 expression and also a higher proportion of phosphorylated eIF4G1 within the eIF4G1 pool. The highest elevation of phosphorylation in eIF4G1 occurs at Ser1035 (**Figure 6F**) and Ser1087 (**Figure 6I**), as the abundances of two phosphor-peptides in tumors are 3.812 and 3.919 times the corresponding abundances in healthy tissues. The sites with significantly elevated phosphorylation in tumors, including Ser1035 (**Figure 6F**), Ser1084 (**Figures 6G–6H**), Ser1087 (**Figure 6I**), Ser1099 (**Figure 6J**), Ser1201 (**Figure 6K**), Ser1216 (**Figure 6L**), Ser1238 (**Figure 6M**) and Ser1245 (**Figure 6N**), are mostly clustered at the interdomain linker region (IDLR) that separates the HEAT-1 and HEAT-2 domains in eIF4G1. This observation dovetails with a previous report of intense phosphorylation of eIF4G1 within its IDLR in proliferating cells during mitosis (Dobrikov et al., 2014). Moreover, the intense phosphorylation at the eIF4G1 IDLR has been shown to enhance the binding of HEAT-2 domain to eIF4A1, which may alter the regulatory strength of eIF4G1 in tumors (Dobrikov et al., 2014).

**Fig. 6.**
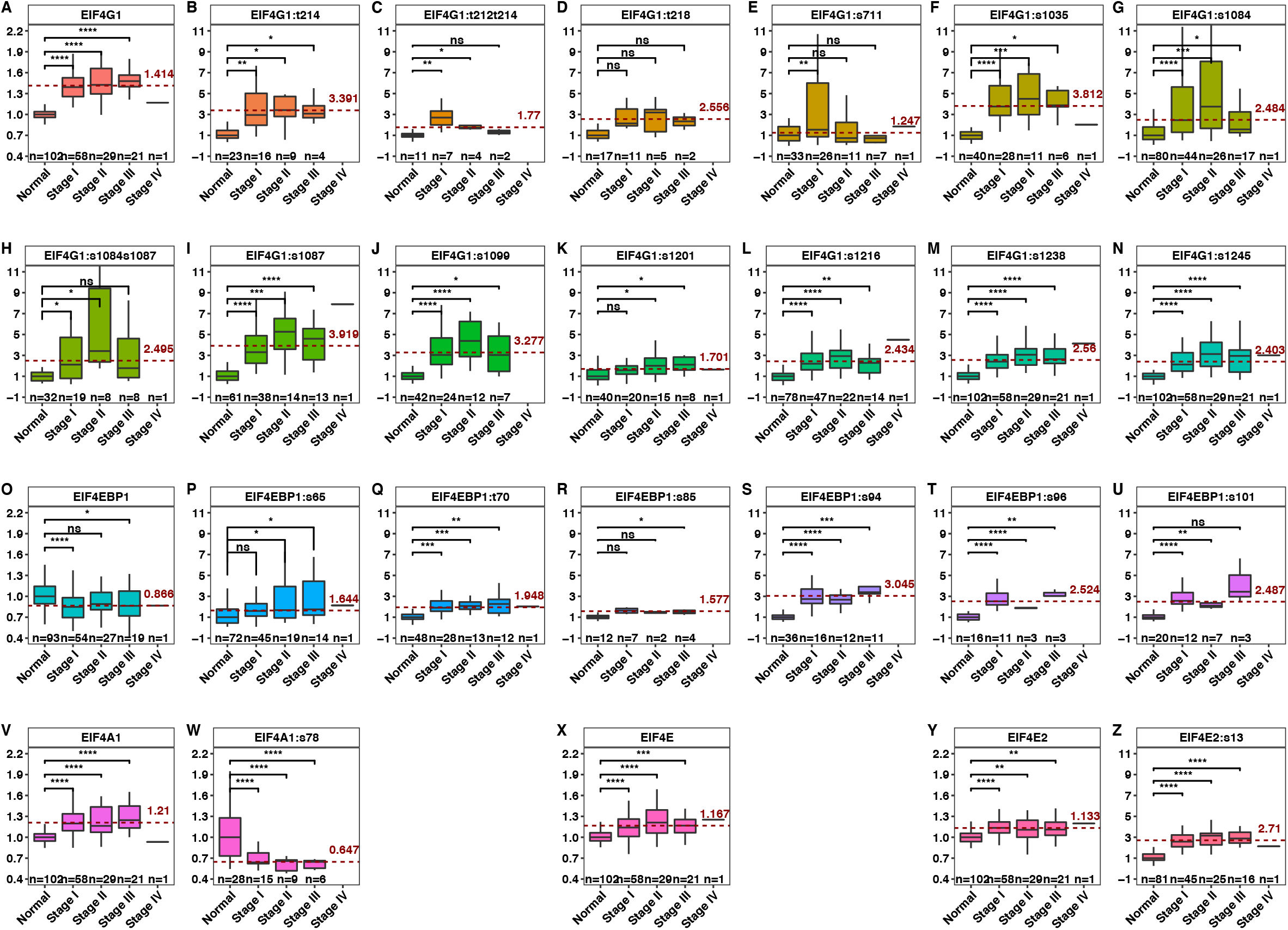
Phosphorylation of a regulatory region in eIF4G1 is elevated in lung adenocarcinoma. (**A**) The relative abundances of eIF4G1 protein from normal lung tissues and different stages of lung adenocarcinomas. The analysis of eIF4G1 protein abundance was performed with the proteomics data collected from 109 lung adenocarcinomas and 102 paired normal adjacent tissue samples in CPTAC lung adenocarcinoma (LUAD) discovery study. Protein was quantitated as the relative abundance of detected peptides from eIF4G1 to reference. Y-axis indicates the relative abundance of eIF4G1 normalized to the median value of its abundance in normal adjacent lung tissues. The red dash line marks the average abundance of eIF4G1 in all stage lung adenocarcinomas, compared to that in normal lung tissues. The two-tailed Student’s t tests were performed. ns, not significant; *P ≤ 0.05; **P ≤ 0.01; ***P ≤ 0.001; ****P ≤ 0.0001 (**B** to **N**) The relative abundances of detected phosphor-peptides from eIF4G1 protein in normal lung tissues and different stages of lung adenocarcinomas. The analysis of phosphor-peptide abundance from eIF4G1 was performed on the phosphor-proteomics data collected from 109 lung adenocarcinomas and 102 paired normal adjacent tissue samples in CPTAC lung adenocarcinoma (LUAD) discovery study. The boxplots show relative abundances of phospho-peptides, which contain one or more fully-localized phosphorylation modifications and are mapped to eIF4G1’s sites. Phosphor-peptides were quantitated as the relative abundance of a detected phosphor-peptide from eIF4G1 in a sample to that in the reference. Y-axis indicates the relative abundance of a phosphor-peptide normalized to the median value of its abundance in normal lung tissues. s stands for serine residue and t stands for threonine residue. The red dash lines mark the average abundances of indicated phosphor-peptides in all stage lung adenocarcinomas, compared to those in normal lung tissues. The two-tailed Student’s t tests were performed. ns, not significant; *P ≤ 0.05; **P ≤ 0.01; ***P ≤ 0.001; ****P ≤ 0.0001 (**O**) The relative abundances of 4E-BP1 protein from normal lung tissues and different stages of lung adenocarcinomas. (**P** to **U**) The relative abundances of detected phosphor-peptides from 4E-BP1 protein in normal lung tissues and different stages of lung adenocarcinomas. (**V**) The relative abundances of eIF4A1 protein from normal lung tissues and different stages of lung adenocarcinomas. (**W**) The relative abundances of detected phosphor-peptides from eIF4A1 protein in normal lung tissues and different stages of lung adenocarcinomas. (**X**) The relative abundances of eIF4E protein from normal lung tissues and different stages of lung adenocarcinomas. No phospho-peptide of eIF4E was reported in the CPTAC LUAD data. (**Y**) The relative abundances of eIF4E2 protein from normal lung tissues and different stages of lung adenocarcinomas. (**Z**) The relative abundances of detected phosphor-peptides from eIF4E2 protein in normal lung tissues and different stages of lung adenocarcinomas.

The average abundance of 4E-BP1 protein in lung adenocarcinomas (regardless of cancer stage) is 86.6% of the average abundance in normal lung tissues (**Figure 6O**). The phosphorylation of 4E-BP1 occurs at several sites, including Ser35, Thr36, Thr37, Thr46, Ser65, Thr70, Ser83, Ser85, Ser86, Ser94, Ser96 and Ser101 (**Figures 6P–6U** and **S12N–S12S**). Our analysis shows significant elevation of 4E-BP1 phosphorylation at Ser65 (**Figure 6P**), Thr70 (**Figure 6Q**), Ser85 (**Figure 6R**), Ser94 (**Figure 6S**), Ser96 (**Figure 6T**) and Ser101 (**Figure 6U**) sites in lung adenocarcinomas, compared to normal adjacent lung tissues. Phosphorylation of Thr37 and Thr46 affects adjacent motif 1 that is responsible for direct eIF4E binding (Sekiyama et al., 2015b); The priming phosphorylation at Thr37 and Thr46 catalyzed by mTOR is required for subsequent phosphorylation at Ser65 and Thr70 (Gingras et al., 1999). When 4E-BP1 is hyperphosphorylated at these four amino-acid residues, it no longer sequesters eIF4E, allowing the binding of eIF4G1 to eIF4E and CDTI (Gingras et al., 1999). However, phosphorylation of Ser65 and Thr70 alone, both of which are within motif 2, a proline–turn–helix segment, is insufficient to block 4E-BP1’s binding to eIF4E (Gingras et al., 2001). We found the phosphor-peptides with Thr37 and Thr36 in only 8 samples (less than 5% of total samples), suggesting a low incidence of Thr37/Thr46–priming phosphorylation (**Figures 12N–12P)**. The abundance of phospho-peptides with Ser65 or Thr70 in tumors is 1.644 and 1.948 times the corresponding abundance in healthy cells (**Figures 6P** and **6Q**), suggesting an increasing proportion of phospho-Ser65 and Thr70 within the total 4E-BP1 pool. The highest tumorspecific elevation of phosphorylation occurs at Ser94, Ser96 and Ser101, as the abundances of phosphor-peptides with those sites in tumors are 3.045, 2.542 and 2.478 times the corresponding abundances in healthy cells (**Figures 6S–6U**). 4E-BP1 phosphorylated at Thr-70, Ser-83, and Ser-101 has been reported to bind to eIF4E during mitosis and not to specifically inhibit translation initiation (Sun et al., 2019). Our results suggest 4E-BP1 phosphorylation is elevated in lung adenocarcinoma, but perhaps is not sufficient to block 4E-BP1’s sequestration of eIF4E.

The average abundance of eIF4A1 protein in all stages of lung adenocarcinomas is 21% above those in normal lung tissues (**Figure 6V**). The phosphorylation of eIF4A1 occurs at Tyr70, Ser78, Ser205 and Thr207 sites (**Figures 6W** and **S12T–S12V**). The average abundance of eIF4A1 phosphorylated at Ser78 in lung adenocarcinomas is 64.7% of that in normal adjacent tissues (**Figures 6W**), suggesting a tumorspecific dephosphorylation of Ser78. Ser78 residue is within the ATP-binding motif (P loop) that directly binds to ATP or ADP, and the negative charge of Ser78 phosphorylation likely interferes with the binding of ATP/ADP that are also negatively charged (Schutz et al., 2010). We reason that Ser78 dephosphorylation is preferred for optimal ATP binding, which may subsequently promote the binding of eIF4A1·ATP to HEAT-1 domain of eIF4G1 (Marintchev et al., 2009).

The average abundance of eIF4E protein in lung adenocarcinomas is merely 16.7% above that in normal lung tissues (**Figure 6X**). No eIF4E phosphorylation is detected. eIF4E2, a homolog of eIF4E that replaces eIF4E in CDTI to synthesize hypoxia-response proteins in tumors (Uniacke et al., 2012), is detected in proteomics data. The average abundance of eIF4E2 in tumors is 13.2% above that in normal lung tissues (**Figure 6Y**). One phosphorylation site of eIF4E2 is found at Ser13 (**Figure 6Z**). The elevation of Ser13 phosphorylation is tumor-specific, as its average tumor versus normal change is 2.71 times. Since Ser13 is located at the N-terminal of eIF4E2 that is distal to the 4E-BP1 and cap-binding domains, it is inconclusive whether its phosphorylation affects CDTI or not. In summary, the phosphorylation statuses of eIF4F subunits in tumors suggest more eIF4A1·eIF4G1 interaction, and more 4E-BP1oeIF4E interaction, in tumors than in healthy tissues.

## DISCUSSION

Cellular mRNA abundance is regulated by transcription and mRNA degradation, whereas protein abundance is largely determined by translation and protein degradation (Schwanhausser et al., 2011). Regarding gene expression, mRNA levels are generally different from protein levels due to the homeostasis of biomolecules. For stable proteins, however, degradation may be discounted and abundance may be estimated as a function of synthesis. The protein synthesis rate generally correlates with corresponding mRNA concentration and translational efficiency (Hershey et al., 2012). Since housekeeping genes such as those used for translation regulation are characterized by both mRNA and protein stability, abundance of one may be used as a proxy for abundance of the other. Furthermore, the subunits of eukaryotic protein complexes tend to be synthesized at rates in proportion to their stoichiometry (Taggart and Li, 2018); hence mRNA ratios and protein ratios may be deemed proportional to each other. In the specific case of *EIF4F* genes, RNA levels provide a good measure for protein levels (Gillette et al., 2020; Mertins et al., 2016). Therefore, quantitative RNA-Seq from pan-cancer types may reasonably be used to infer the abundance of eIF4F protein components in tumors; and our analysis is predicated on such inference.

### Positive selection of *EIF4G1* in cancers

The constituent types of somatic copy number variations (CNVs) are gene amplification, duplication, and heterozygous and homozygous deletion (Heitzer et al., 2016). CNVs, common among genetic alterations, are detected in 90% of solid tumors. They occur stochastically as a consequence of errors in DNA repair of double-strand breaks by the breakage-fusion-bridge mechanism (Coquelle et al., 1997; Mc, 1951). In the wake of breakage, whether cancer cells select for amplifications in some settings or copy number loss in others is often influenced by the nature of genes that give a survival advantage to a clone in its microenvironment (Heitzer et al., 2016). *EIF4G1, EIF4E* and *EIF4EBP1* locate at the chromosome region 3q27.1, 4q23 and 8p11.23 respectively. These three regions have been previously reported to be susceptible to chromosomal breaks. *EIF4G1* amplification/duplication may be related to the chromosomal rearrangements at 3q27 that occurs in up to 45% of diffuse large B-cell lymphoma (DLBC) (Lo Coco et al., 1994). Truncation and amplification/duplication of other genes including *BCL6* has been observed to associate with the chromosomal aberrations of 3q27 in DLBC (Huang et al., 2019; Ye et al., 1993). *EIF4E* deletion may be related to the chromosomal break at 4q23, a common chromosomal fragile site that is susceptible to frequent gaps or breaks in response to replication stress, as reported in bone marrow cells (Morgan et al., 1988). The 8p11-p12 chromosomal region containing *EIF4EBP1* is a frequent target for amplification and deletion in breast cancer (Adelaide et al., 1998; Gelsi-Boyer et al., 2005), as well as translocation in hematologic malignancies such as myeloproliferative disorder (Macdonald et al., 1994) and acute monocytic leukemia (Lai et al., 1992). Therefore, the chromosomal locations of *EIF4F* genes trigger the high incidences of their CNVs.

Because multi-protein complexes depend upon the stoichiometry of their subunits to perform biological functions, CNVs of protein complex genes can have exaggerated effect and are therefore less common than CNVs of other genes (Papp et al., 2003; Schuster-Bockler et al., 2010). However, we found that 7.74% of all tumors contain *EIF4G1* amplification and 26.79% contain duplication (**Figure 1C**), that *EIF4G1* CNV is associated with *EIF4G1* mRNA overexpression (**Figure 1E**), and that elevated *EIF4G1* expression correlates with poor survival in cancer patients (**Figure 4A**). These observations provide evidence for strong positive selection by tumors towards increased effect of *EIF4G1*, implying its tumor-beneficial function during cancer evolution (Bakhoum and Landau, 2017; Zack et al., 2013). In contrast, we found that 30.21% of all tumors contain *EIF4E* heterozygous deletion (**Figure 1C**), yet nonetheless *EIF4E* expression is marginally elevated in primary tumors compared to healthy adjacent tissues (**Figure S3C**). It may be, therefore, that tumors with *EIF4E* heterozygous deletion upregulate *EIF4E* expression to a level similar to healthy tissues with intact *EIF4E* genes, probably via alternative regulation such as transcription. Those findings echo a previous report that mice with heterozygous deletion of *EIF4E* (*EIF4E*^+/-^) have normal development and global protein synthesis (Truitt et al., 2015). That study also shows that *EIF4E*^+/-^ mice have impeded KRas-induced lung tumorigenesis. Based on that evidence, we perceive two possible explanations for our data, neither of which excludes the other: 1) *EIF4G1* overexpression to some extent compensates for *EIF4E* activity in tumors; and 2) tumors well tolerate *EIF4E* heterozygous deletion. The dramatic difference in CNV statuses of *EIF4G1* and *EIF4E* (**Figure 1D**), both occurring at high incidence, suggest a natural selection process by which tumors adjust the functional impact of those two subunits.

Physiological stress within the tumor microenvironments may apply selective pressures that favor overabundance of *EIF4G1*. For example, hypoxia increases generation of reactive oxygen species (ROS) (Chandel et al., 1998; Fruehauf and Meyskens, 2007), so efficient translation of ROS response genes is necessary for cancer cell survival. ROS response genes, such as the ferritin gene (*FTH1*), can be regulated by CDTI, as fewer ferritin proteins are translated in *EIF4E*^+/-^ murine embryonic fibroblasts (MEFs) than in wildtype MEFs after transformation (Truitt et al., 2015). *FTH1* also contains IRES at its 5’UTR and are reportedly subject to CITI regulation via eIF4G1 in human lung cancer cell lines (Daba et al., 2012). Human breast cancer cells under radiation treatment rely on eIF4G1 overexpression and CITI to translate genes for cell survival and DNA repair (Badura et al., 2012). Therefore, under physiological conditions, tumors may be pressured to select for *EIF4G1* over *EIF4E* and may then rely upon CITI to survive.

### Computational analyses of *EIF4F* expression conform to biochemical findings

Our computational analysis results complement extant biochemistry studies. For example, eIF4G1 contains two eIF4A1 binding domains: the HEAT1 domain at the middle portion of eIF4G; and the HEAT2 domain at the C-terminal portion, which itself contains a separate HEAT3 domain binding to the eIF4E kinases Mnk1 and Mnk2. eIF4G1 can take a folded conformation that allows both HEAT1 and HEAT2 domains to bind to a single eIF4A1 using IDLR as a hinge (Marintchev et al., 2009). eIF4G1 can also take an elongated conformation that allows HEAT1 and HEAT2 each to bind to its own distinct eIF4A1 (Nielsen et al., 2011). Assembly of eIF4A1·eIF4G1 complexes in either 1:1 or 2:1 ratios has been reported by biochemical reconstitution studies (Korneeva et al., 2001; Nielsen et al., 2011). In accordance with this notion, we found a 1:1 ratio between *EIF4A1* and *EIF4G1* mRNA abundance in most healthy tissues (**Figures S3D and S4B**), and a more evident 2:1 ratio in some cancer types and metastatic tumors (**Figures 2B and S3D**). Moreover, binding of HEAT-1 to eIF4A is essential for translation initiation, whereas binding of HEAT-2 only modulates eIF4A1 activation (Morino et al., 2000). We observed significant dephosphorylation of eIF4A1 at S78 (**Figures 6W**) in lung adenocarcinoma that likely facilitates its ATP-binding and favors an overall closed conformation of eIF4A, optimal for HEAT-1 binding (Marintchev et al., 2009). We also observed cancer-specific hyperphosphorylation of eIF4G1 (**Figures 6F–6N)** at IDLR, which has been shown to promote eIF4A1’s binding to the HEAT2 domain during mitosis (Dobrikov et al., 2014). Those findings suggest that tumors prefer interaction between eIF4G1 and eIFA1 via both HEAT-1 and HEAT-2 domains to enhance translation initiation. The stoichiometric shift of *EIF4A1*:*EIF4G1* and altered phosphorylation statuses of eIF4G1 and eIF4A1 in tumors may imply a more favorable 2:1 ratio for eIF4A1·eIF4G1 complex.

Downregulation of 4E-BP1 – and consequent increase of eIF4E availability, eIF4E:4E-BP1 ratio, and CDTI – have been reported to counter the action of mTOR inhibitors in transformed MEFs (Alain et al., 2012). Consistently, our RNA-seq analysis show that *EIF4E*:*EIF4EBP1* ratios are much higher than 1 in healthy tissues (**Figure S4B**), which suggests that that healthy tissues tend to increase eIF4E:4E-BP1 ratio and eIF4E availability to ensure CDTI. 4E-BP1 overexpression has been reported in many cancer types (Musa et al., 2016). In human breast cancer cells, 4E-BP1 overexpression leads to the inhibition of CDTI, and enhances translation of mRNAs required for growth under hypoxic conditions using CITI (Braunstein et al., 2007). Consistently, we found that *EIF4E*:*EIF4EBP1* ratios are close to 1 in most cancer types (**Figure 2B**), suggesting that tumors tend to decrease eIF4E:4E-BP1 ratio and eIF4E availability. Furthermore, the low incidence of Thr37/Thr46 phosphorylation of 4E-BP1 in lung adenocarcinoma suggests that physiologically stressful conditions intrinsically inhibit mTOR and capacitate binding of 4E-BP1 to eIF4E, with consequent inhibition of CDTI (**Figures S12O** and **S12P**). Those observations seem to explain a lack of significant association of *EIF4E* and *MTOR* expression with poor prognosis in all TCGA or in lung adenocarcinoma patients. The change of *EIF4E*:*EIF4EBP1* ratio and phosphorylation status of 4E-BP1 in tumors may imply a 4E-BP1·eIF4E complex formation, a potential hindrance for CDTI.

Phosphorylation of eIF4E serine 209 reportedly has been detected in most malignant melanomas with constitutively active *BRAF^V600E^* mutation and associated with reduced survival in patients (Boussemart et al., 2014; Carter et al., 2016), since Mnk1/Mnk2 are directly stimulated by Erk or p38 from the Ras/Raf/Mek pathway (Waskiewicz et al., 1997). In addition, eIF4E is phosphorylated only as a part of the eIF4F complex, when Mnk1 is recruited directly via the HEAT3 of eIF4G1 (Pyronnet et al., 1999). eIF4E phosphorylation has been reported in 39.9% of biopsies from non-small cell lung cancer predominantly with active Akt (Yoshizawa et al., 2010). Mechanistically, phosphorylation of 4E-BP1 by PI3K/Akt/mTOR1 allows eIF4E to bind to eIF4G1. However, active Ras/Raf/Mek and PI3K/Akt/mTORC1 pathways do not guarantee eIF4E phosphorylation. For example, phosphorylation of eIF4G1 IDLR promotes binding of Mnk1 to HEAT3 (Dobrikov et al., 2014). Yet, despite Mnk1 activation, eIF4E has been observed to remain unphosphorylated in mitotic cells, due to hyperphosphorylation of eIF4G and hypophosphorylation of 4E-BP1, both of which diminish the binding of eIF4E to eIF4G (Pyronnet et al., 2001). As seen in **Figure 6**, hyperphosphorylation of eIF4G1 and hypophosphorylation eIF4E from CPTAC lung adenocarcinomas – which for the most part are primary tumors harboring *EGFR* or *KRAS* mutation (Gillette et al., 2020) – seem to confirm an indispensable role of eIF4G1 to facilitate eIF4E phosphorylation. Although a previous study, based on computational predictions, suggested that the eIF4G1 IDLR could influence eIF4E·eIF4G1 via intramolecular interaction by forming antiparallel β-hairpin with the eIF4E-binding motif, which might be regulated by eIF4G1 phosphorylation at IDLR (Dobrikov et al., 2018), no detailed mechanism has yet been confirmed by protein structure study.

Stoichiometry measurements in mammalian eIF4F varied greatly among complexes purified by different biochemical approaches. Nonetheless, as the limiting factor for CDTI (Haghighat and Sonenberg, 1997), eIF4E abundance was reported much lower than other eIF4F components (Duncan et al., 1987; Galicia-Vazquez et al., 2012), Consistently, our RNA-Seq analysis suggested that *EIF4G1* and *EIF4A1* expressions are much higher than *EIF4E*. *EIF4E*, *EIF4A1* and *EIF4G1* promoters all contain the E-box motifs (5’CACGTG3’) directly regulated by c-Myc via transcription (Jones et al., 1996; Lin et al., 2008), which enables feedforward regulation between c-Myc and eIF4F (Komili and Silver, 2008). The constant 1:1 ratio for *EIF4G1*:*EIF4A1* and 8:1 for *EIF4G1*:*EIF4E* in healthy tissues confirm a previous report that transcription of *EIF4F* subunits for CDTI is highly coordinated (Lin et al., 2008). Furthermore, the *EIF4E* promoter contains hypoxia-responsive elements (HREs) that can be directly regulated by HIF1α under hypoxic conditions (Azar et al., 2013; Yi et al., 2013). Since HREs overlap the E-box motifs in *EIF4E* promoter, c-Myc and HIF1α compete for the transcription regulation of *EIF4E* (Gordan et al., 2007), which may perturb the coordinated transcription between *EIF4G1* and *EIF4E*. Consistently, we observed no constant ratio of *EIF4G1*:*EIF4E* in most cancer types. The *EIF4EBP1* promoter also contains HREs that are regulated by HIF1α in cooperation with SMAD4 under hypoxia condition (Azar et al., 2013). The 4:1 ratio of *EIF4G1*:(*E+4EBP1*) in tumors (**Figure 2D**) probably reflects the interplay among c-Myc, HIF1α and SMAD4 on transcription regulation. Additional biochemical investigation will be needed to uncover the detailed mechanism.

### CITI is one manifestation of dysregulated translation initiation in cancers

Elevated translation initiation activity is regarded as the major evidence of translation dysregulation in malignant tumors (Ruggero, 2013), particularly because eIF4E overexpression has been reported in many cancer types including prostate, breast, stomach, colon, lung, skin, and the hematopoietic system (Graff et al., 2009; Hsieh and Ruggero, 2010; Rosenwald et al., 1999). Our PCA data confirmed that *EIF4E* expression is a key distinguishing factor between tumor samples and healthy tissues (**Figure 3A**), even though the level of *EIF4E* overexpression in tumors is small over healthy tissues due to the frequent heterogenous deletion (**Figure S3C**). In addition, our findings showed that a combination of seven *EIF4F* genes adequately differentiate malignant tumors from healthy tissues, regardless of cancer types or tissue types. *EIF4F* genes may serve as potential diagnostic biomarkers for identifying patients with malignant conditions using minimally invasive measurement of circulating DNA or RNA. Furthermore, the expression variability of *EIF4F* is well able to distinguish healthy tissues from each other by tissue type (**Figure 3D**), but is much less able to do the same for tumors (**Figure 3E**). This observation strongly supports the notion that eIF4F activities are distinctly-regulated in individual healthy tissue types to carry out tissue-specific functions, and such regimes are lost across cancer types (Ruggero, 2013).

Our work reveals additional aspects of eIF4F dysregulation in cancer by showing that individual *EIF4F* subunits vary in their prognostic strengths. *EIF4E* is a weak oncogene in its own right, because in the absence of other factors its overexpression induces cellular senescence (Ruggero et al., 2004). When both *EIF4E* and *MYC* are overexpressed, however, they cooperatively promote B-cell lymphomagenesis in a murine lymphoma model, because c-Myc overrides cellular senescence in MEFs, whereas eIF4E reduces apoptosis (Ruggero et al., 2004). Consistently, *EIF4E* expression alone fails to predict poor prognosis in all TCGA cancer patients with three survival analyses (**Figure 4**). However, *EIF4G1* and *EIF4A1* expressions each show strong correlation with poor prognosis in KM analyses, suggesting their crucial functions in malignancy. In our joint survival impact assessments of initiation factors from multivariate Cox analysis, *EIF4G1* but not *EIF4E* or *EIF4A1* expression significantly contributes to poor prognosis in all tumor cases combined, and also in lung tumor cases specifically (**Figures 4G** and **4H**). By implication, the disease-promoting attribute of *EIF4A1* may depend on *EIF4G1* in tumors, which is in line with a prior biochemical finding that mRNA-binding of eIF4A1 depends on eIF4G1 (Marintchev et al., 2009). Although correlation does not guarantee causal relation, our findings raise the possibility that *EIF4G1* controls the translation initiation for malignancy progression. This observation suggests that eIF4E function is particularly dysregulated in cancer.

A dramatic phenotype of eIF4F dysregulation is functional separation between eIF4E and eIF4G1/eIF4A1 to control different mRNA targets, which has been reported in breast cancer and melanoma (Badura et al., 2012; Joyce et al., 2017). We found that *EIF4G1, EIF4A1* and *EIF4E* equally and strongly correlate with numerous genes associated with house-keeping functions, particularly in healthy tissues (**Figures 5C and 5G**). Although correlation analysis with RNA-Seq data does not discern between direct (translational) and indirect (transcriptional) regulatory mechanisms, the results reflect highly similar functions of *EIF4A1, EIF4G1* and *EIF4E*, and are in line with biochemical findings that the majority of mRNA translation in healthy tissues rely upon the canonical eIF4E·eIF4G·eIF4A1 complex for CDTI (Kwan and Thompson, 2019). The same posCORs for *EIF4F* genes from healthy tissues have much higher correlation coefficients than those from tumors, providing strong evidence for functional separation as a prevailing phenomenon for eIF4F dysregulation in cancers.

Our work reveals that eIF4F dysregulation is rooted in the diverse translation initiation mechanisms among mRNA templates. For example, CDTI via the canonical eIF4G1·eIF4A1·eIF4E complex is suppressed by 4E-BP1 during mitosis (Badura et al., 2012; Joyce et al., 2017), wheras CITI by IRES binding is enhanced (Badura et al., 2012). Strikingly, we discovered in lung adenocarcinoma, *EIF4G1, EIF4A1* and *EIF4EBP1*, but not *EIF4E*, preferentially correlate with genes involved in pathological progression and mitotic regulation (cluster 1 genes **Figures 5G** and **5H**), which suggests an intriguing scenario: eIF4G1 in malignant tissues may have been drafted to regulate CITI for genes that are critical for cancer cell survival. Our data further suggest that the formation of a variant eIF4G1·eIF4A1 complex for CITI likely involves 4E-BP1, consistent with a former report that 4E-BP1 promotes the switch from CDTI to CITI for survival genes in human breast cancer cells (Braunstein et al., 2007). Thus, tumors impose requirements of both CDTI and CITI among diverse mRNA templates for survival benefit.

## Conclusion

Overall, our work suggests that despite an apparent tendency to dysregulate translation machineries of protein synthesis, tumors seem to persist in employing eIF4G1 for translation initiation. Cancers selectively overexpress *EIF4G1* to achieve stoichiometric ratios with *EIF4E* and *EIF4A1* that differ from mRNA ratios typical of the trimetric eIF4F complex in healthy cells. These stoichiometric changes in mRNAs may amount to a “signature” that permits use of *EIF4F* and relevant genes as biomarkers for malignancy and may also support the use of *EIF4G1* as a prognostic indicator in cancer patients. *EIF4G1, EIF4A1* and *EIF4E* equally and strongly correlate with genes associated with house-keeping functions, particularly in healthy tissues. In lung tumors, *EIF4G1* and *EIF4A1*, but not *EIF4E*, each preferentially correlate with a group of genes involved in pathological progression and cell proliferation. In lung tumors, a lack of phosphorylation of 4EBP1 at Thr37/Thr46 leaves it free to inhibit eIF4E and CDTI, and hyperphosphorylations at eIF4G1’s IDLR promote its binding to eIF4A1, both of which likely contribute to the reassignment of eIF4G1 to CITI.

Our work indicates that a prevailing regulatory role of eIF4G1 in translation initiation defines malignant phenotype and influences disease progression, in most cancers. Tumors dysregulate biological “housekeeping” pathways that are usually regulated by cap-dependent translation initiation in healthy tissue, yet still strengthen regulation of cancer-specific pathways in cap-independent contexts. Our work further suggests that lung adenocarcinomas might alternate between cap-dependent and -independent translation initiation mechanisms to sustain optimal cellular functions, through eIF4G1’s adaptable biochemical interactions with eIF4F subunits. These findings reveal clinical relevance of assorted interactions among eIF4F components, offer a means of ranking the susceptibility of cancer types to eIF4F inhibition, and highlight the value of computational analysis to validate and guide biochemical research efforts in the hunt for new and better cancer treatments.

## SUPPLEMENTARY FIGURE LEGENDS

**Fig. S1.**
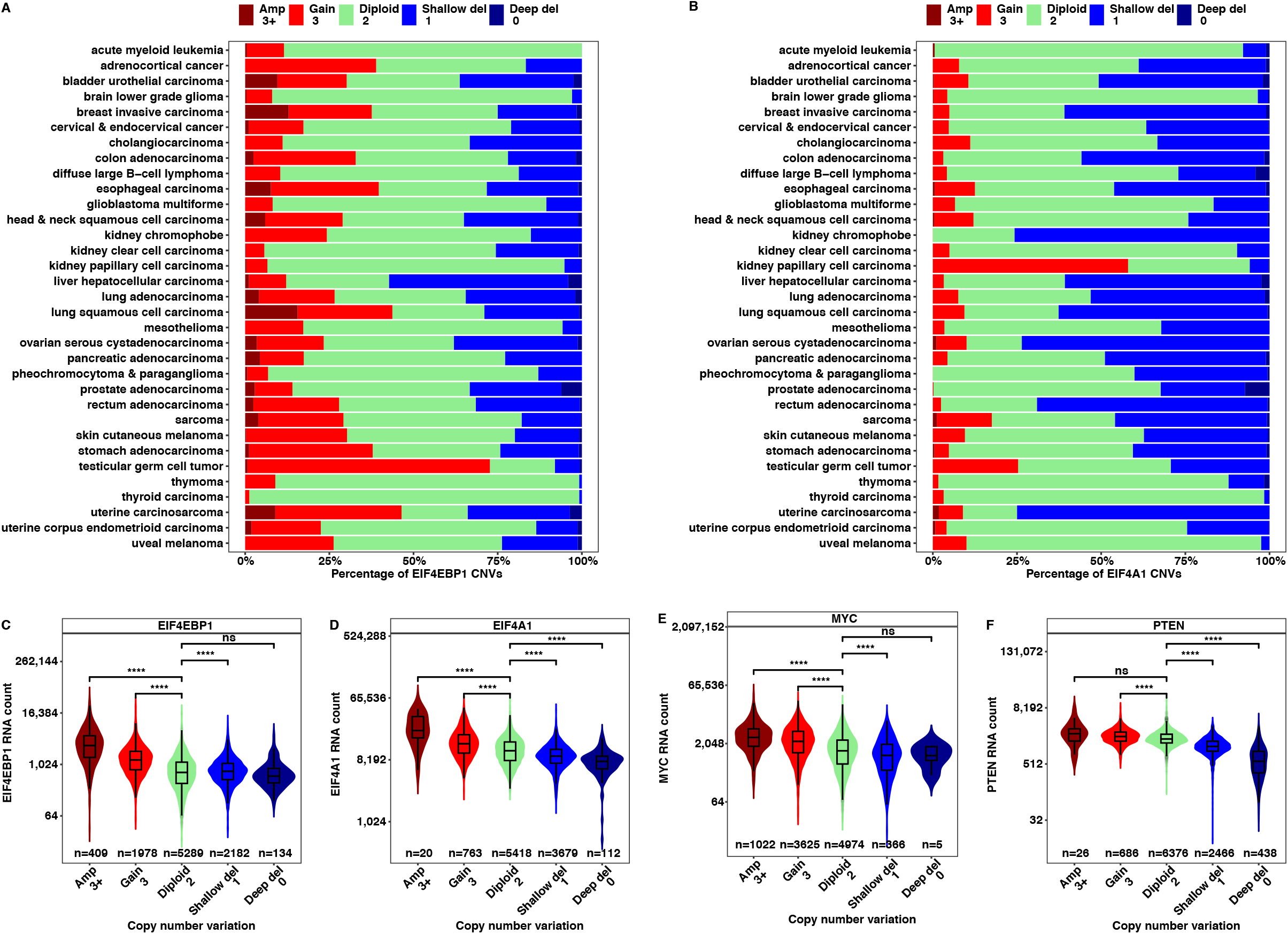
Copy number variations of *EIF4EBP1* and *EIF4A1* in malignant tumors. (**A** and **B**) The CNVs were estimated by GISTIC2 method using the whole genome microarray data. The estimated CNV values were categorized into five group with the following thresholds: 0, 1, 2, 3, 3+. These five groups represent deep deletion (possibly homozygous deletion), shallow deletion (possibly heterozygous deletion), diploid, low-level copy number gain (duplication), or high-level copy number gain (amplification) respectively. The stacked bar plots show the percentages of CNVs for *EIF4EBP1* and *EIF4A1* in different TCGA tumor types. (**C** to **F**) The associations between RNA expression and CNV statuses were analyzed for *EIF4EBP1 EIF4EA1, MYC* (positive control) and *PTEN* (negative control) in all TCGA tumor biopsies combined. The violin plots show the median values of RNA-Seq counts (transcripts per million) of *EIF4G1* and *EIF4EBP1* in the samples with different CNV statuses. The two-tailed Student’s t tests were performed. ns, not significant; *P ≤ 0.05; **P ≤ 0.01; ***P ≤ 0.001; ****P ≤ 0.0001.

**Fig. S2.**
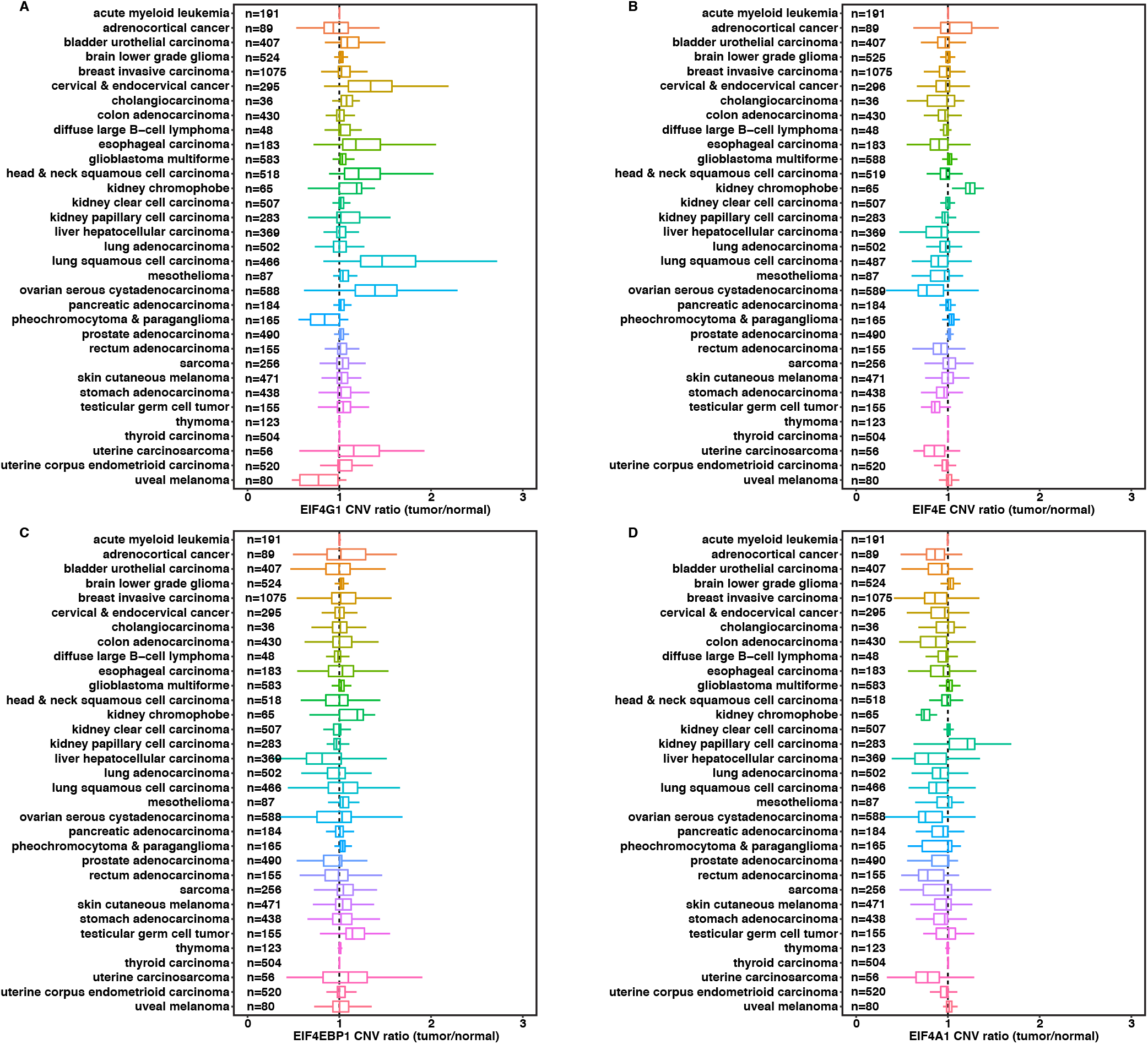
Copy number ratios of *EIF4G1, EIF4EBP1, EIF4E* and *EIF4A1* between malignant tumors and normal tissues. (**A** to **D**) copy number ratios (tumor/normal) were calculated by dividing the copy numbers of *EIF4G1, EIF4E, EIF4EBP1*, and *EIF4A1* in each malignant tumor to the average copy number in adjacent normal tissues, using whole genome microarray data from 33 TCGA cancer groups. The box and whisker plots represent the average CNV ratios of *EIF4E, EIF4G1, EIF4A1* and *EIF4EBP1* across 33 TCGA cancer types.

**Fig. S3.**
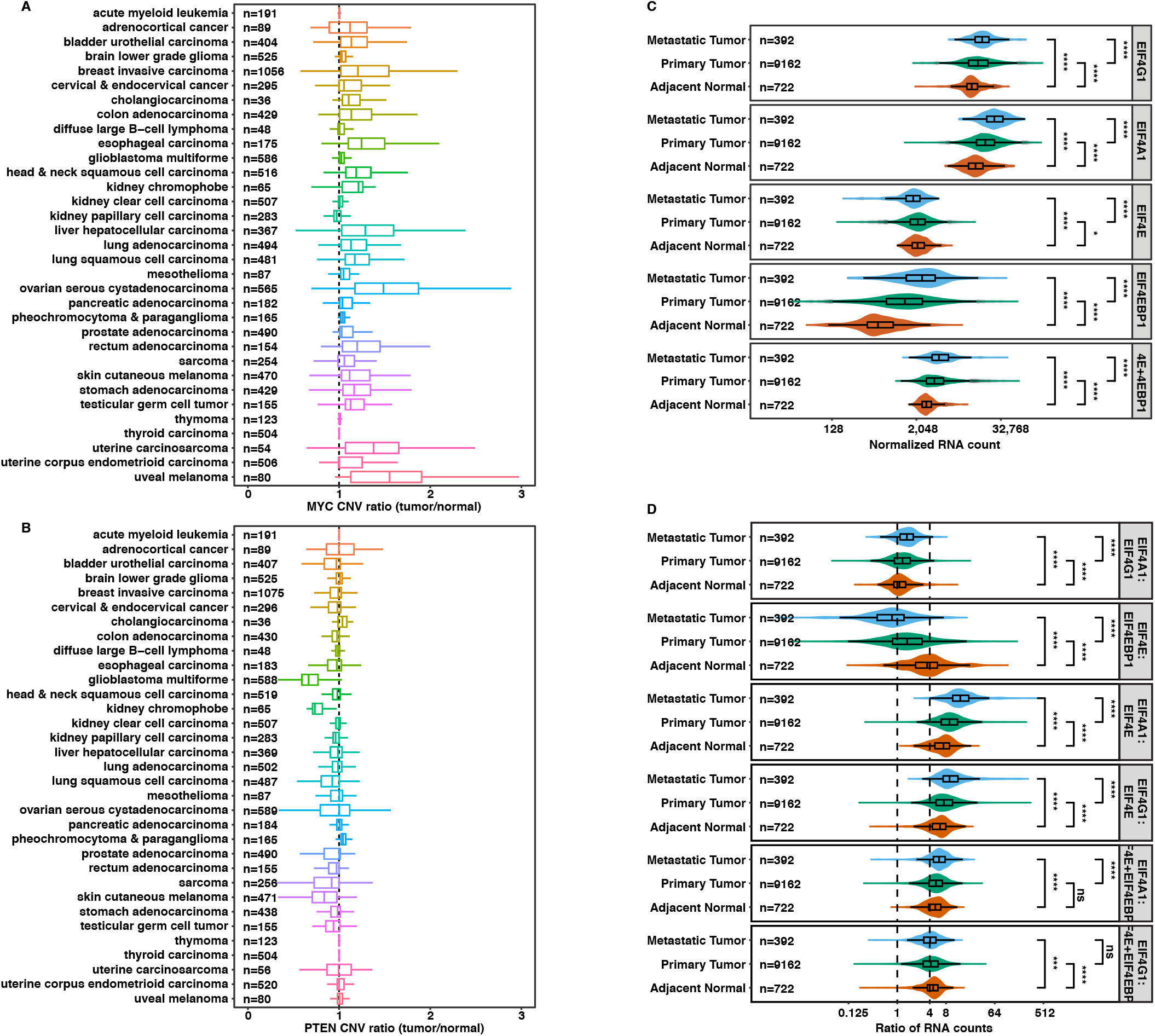
Relative expressions of *EIF4G1, EIF4EBP1, EIF4E* and *EIF4A1* in malignant tumors and normal tissues. (**A** and **B**) a copy number ratio (tumor/normal) was calculated by dividing the copy numbers of *MYC* and *PTEN* in each malignant tumor to the average copy number in adjacent normal tissues, using whole genome microarray data from 33 TCGA cancer groups. The box and whisker plots represent the average CNV ratios of *MYC* and *PTEN* across different cancer types. (**C**) The boxplot shows the average mRNA reads of *EIF4E, EIF4G1, EIF4A1, EIF4EBP1*, and the sum of *EIF4E* and *EIF4EBP1* (*E+EBP1*) in all TCGA cancer patient samples combined. Gene expression was compared among metastatic tumors, primary tumors and adjacent normal tissues. (**D**) The boxplot shows the average ratios between *EIF4F* subunit mRNAs in metastatic tumors, primary tumors and adjacent normal tissues from all TCGA cancer patient samples.

**Fig. S4.**
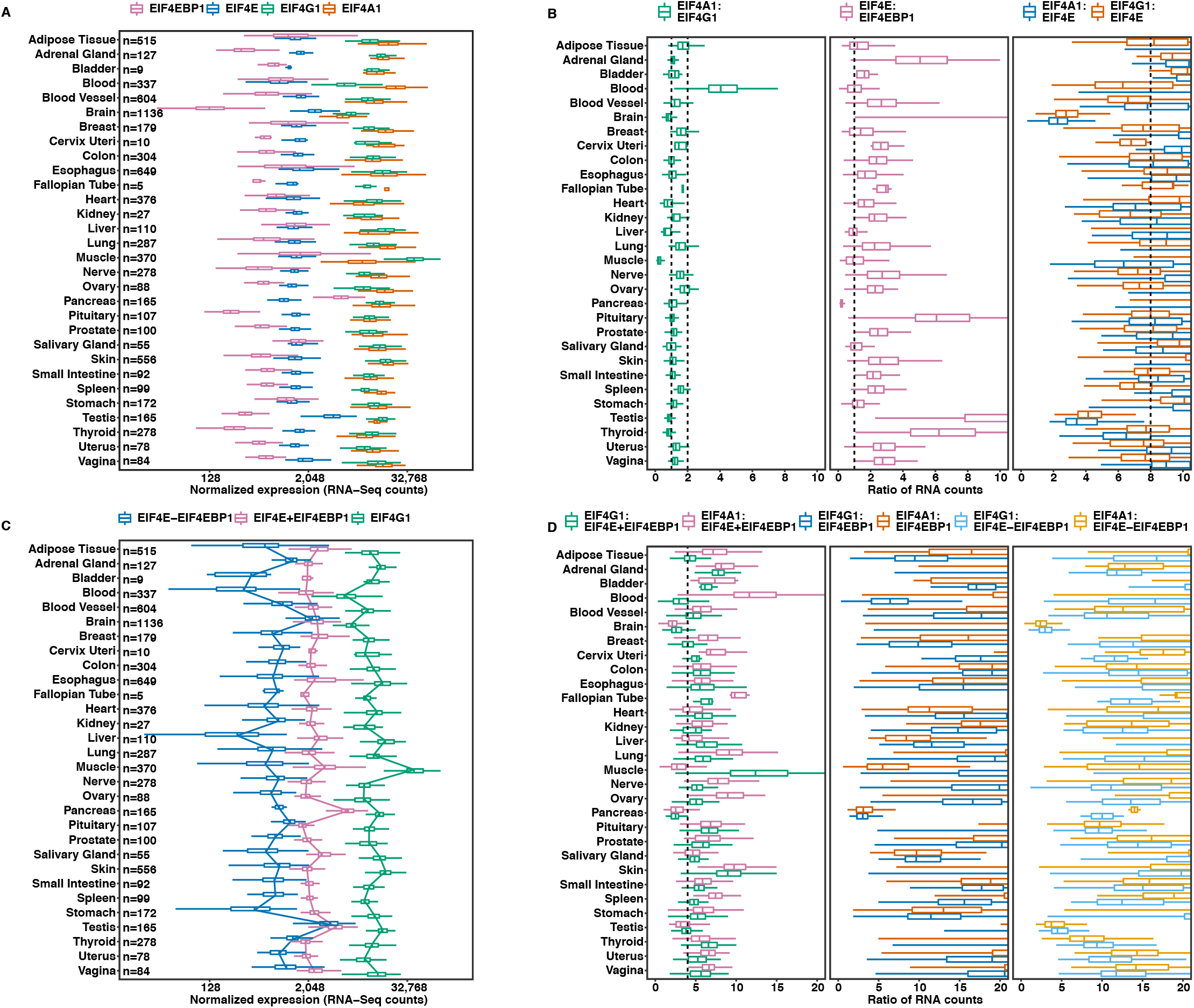
Expressions of *EIF4G1, EIF4EBP1, EIF4E* and *EIF4A1* in healthy tissues. (**A**) The expressions of *EIF4E, EIF4G1, EIF4A1* and *EIF4EBP1* were compared across different healthy tissues using the RNA-Seq data (normalized for batch effects) from 7,388 healthy tissues in Genotype-Tissue Expression (GTEx). (**B**) The ratios of RNA expression were calculated in each tissue samples and compared across healthy tissue types. The dash lines mark 1:1 and 2:1 ratio in the *EIF4A1:EIF4G1* panel, 1:1 ratio in the *EIF4EBP1:EIF4E* panel, and 8:1 ratio in the *EIF4G1:EIF4E* panel. (**C**) The sum expression of *EIF4E* and *EIF4EBP1* (*E+EBP1*), and the difference expression between *EIF4E* and *EIF4EBP1* (*E-EBP1*) were calculated in each healthy tissue sample. They were compared to *EIF4G1* expression across different tissue types. The trendline connects the median values of boxplot. (**D**) the ratios between *EIF4G1* and *E+EBP1*, and the ratio between *EIF4G1* and *E-EBP1* were calculated within each sample and compared across different healthy tissue types. The dash line marks 4:1 ratio in the first panel.

**Fig. S5.**
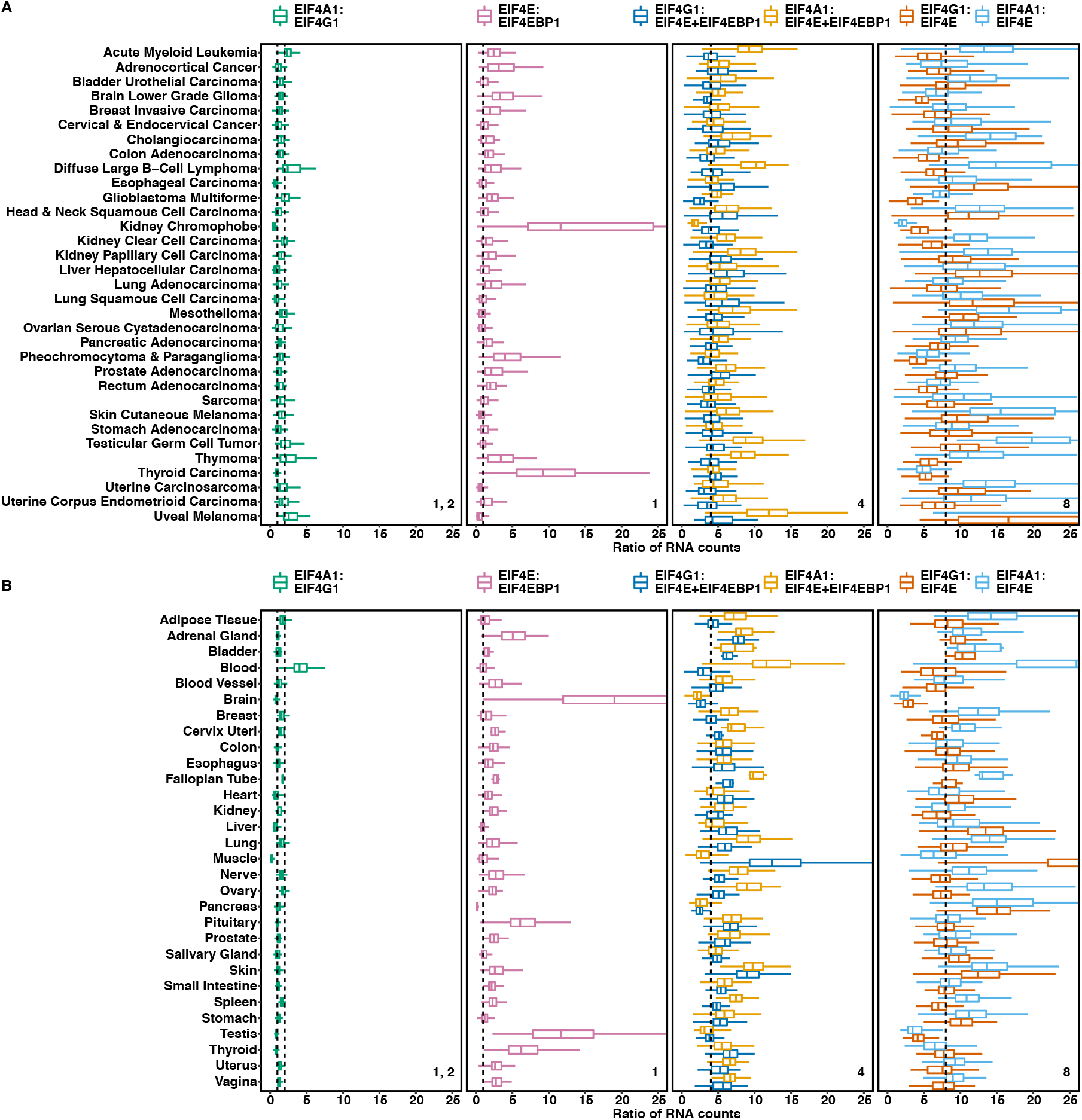
Tumors have unique stoichiometries of *EIF4G1:EIF4A1* and *EIF4G1*:(*EIF4E* + *EIF4EBP1*). (**A** and **B**) the ratios of *EIF4A1:EIF4G1, EIF4E:EIF4EBP1*, *EIF4G1:E+EBP1*, *EIF4A1:E+EBP1, EIF4G1:EIF4E*, and *EIF4A1:EIF4E* were calculated in each tissue samples and compared across TCGA cancer types (**A**) or healthy tissue types (**B**). The dashed lines mark 1:1 ratio in the first two panels for *EIF4A1:EIF4G1, EIF4E:EIF4EBP1*, 4:1 ratio in the third panel for *EIF4G1:E+EBP1* and *EIF4A1:E+EBP1*, and 1:1 ratio in the fourth panel for *EIF4A1:EIF4E* and *EIF4A1:EIF4E*.

**Fig. S6.**
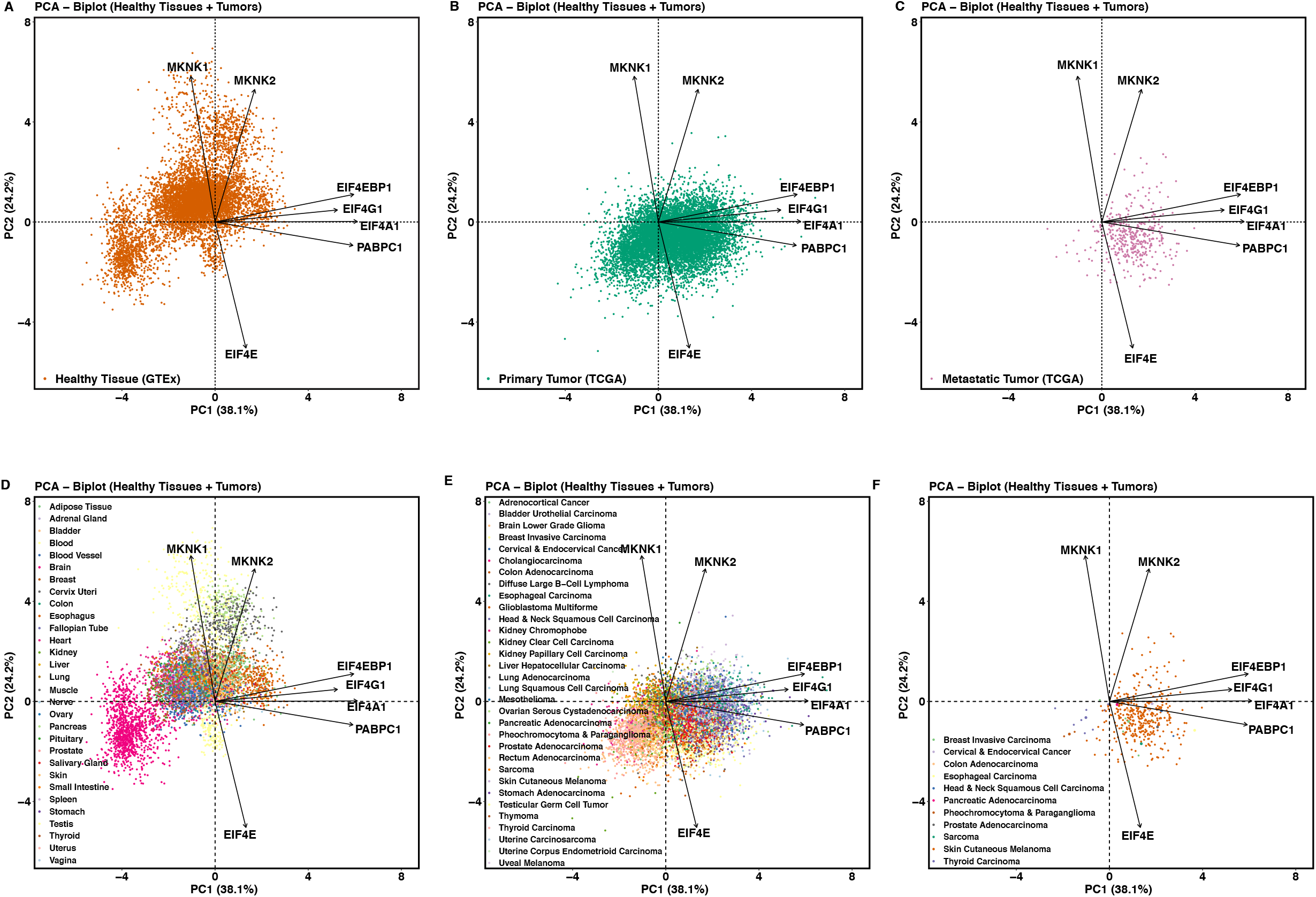
*EIF4G1* and *EIF4E* expressions distinguish tumors from healthy tissues. (**A** and **D**) The PCA biplots show only the healthy tissue samples from the same PCA biplot in **Figure 3A.** The tissue types were not used to construct PCs and individual samples were colored by their tissue types afterwards on the PCA plot (**D**). (**B** and **E**) The PCA biplots show only the primary tumor samples from the same PCA biplot in **Figure 3A.** The cancer types were not used to construct PCs and individual samples were colored by their cancer types afterwards on the PCA plot (**E**). (**C** and **F**) The PCA biplots show only the metastatic tumor samples from the same PCA biplot in **Figure 3A.** The cancer types were not used to construct PCs and individual samples were colored by their cancer types afterwards on the PCA plot (**F**).

**Fig. S7.**
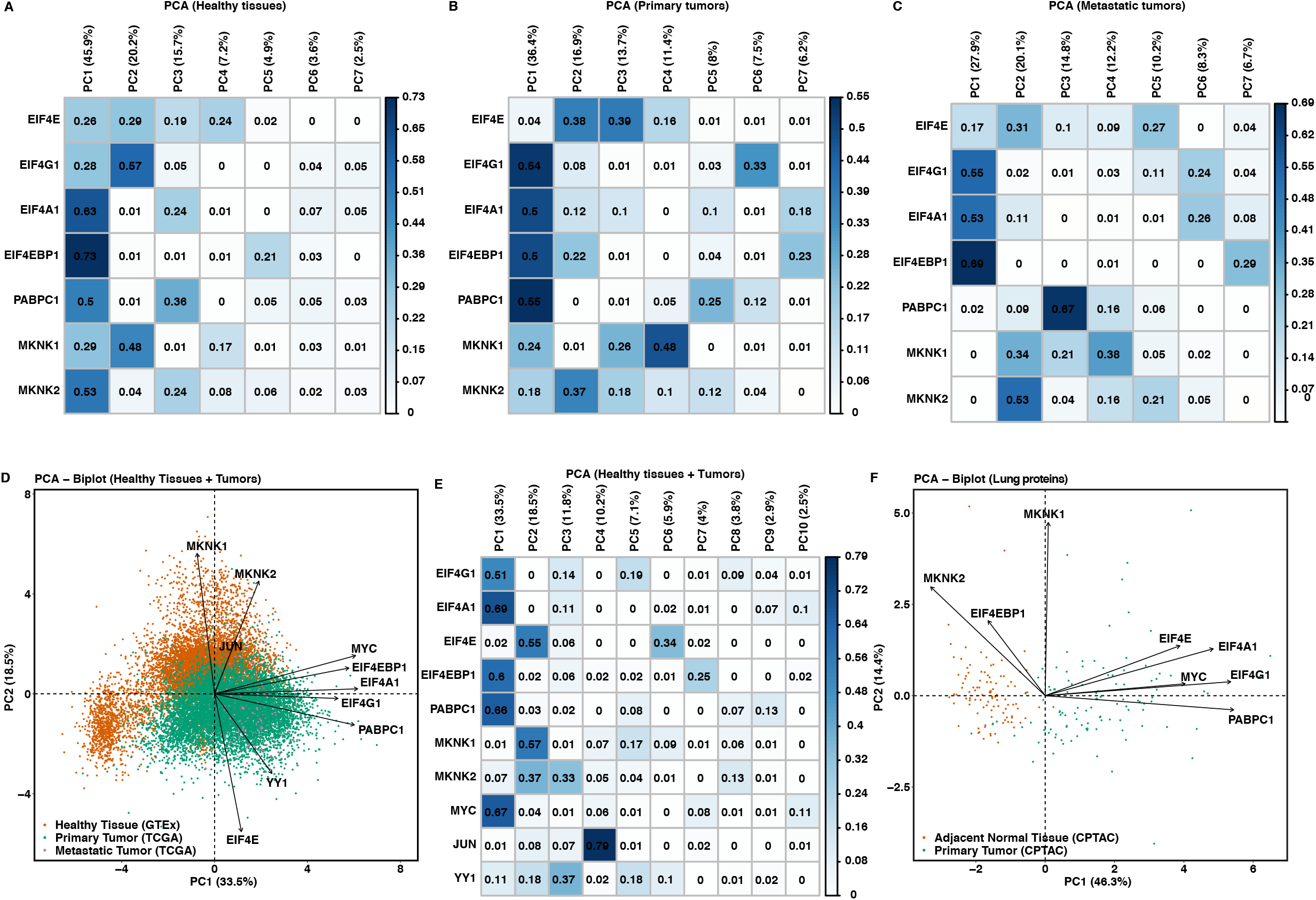
Malignant tumor types are less distinguished from each other by expression of *EIF4F*, in contrast to healthy tissue types. (**A**) The correlation matrix plot for **Figure 3D** shows the percent of variance in which each gene variable is explained by the PC – the values of squared cosines. Cosines refer to the correlation coefficients between the gene variables and PCs. The squared cosine reflects the representation quality of a gene variable on a PCA axis. (**B**) The correlation matrix plot for **Figure 3E** shows the percent of variance in which each gene variable is explained by the PC – the values of squared cosines. (**C**) The correlation matrix plot for **Figure 3F** shows the percent of variance in which each gene variable is explained by the PC – the values of squared cosines. (**D**) The PCA analysis using the standardized RNA expression of *EIF4G1, EIF4A1, EIF4E, EIF4EBP1, PABPC1, MKNK1, MKNK2, MYC, YY1* and *JUN* from all TCGA tumor samples and GTEx normal tissue samples. The PCA biplot shows the first two PCs along which the samples vary the most. Axis title shows the percentage of variances (squared loadings) explained by PC1 or PC2. Axis values show the PCA scores of individual samples. Arrows show the influence of each gene variable on the PCs. (**E**) The correlation matrix plot for (**D**) shows the percent of variance in which each gene variable is explained by the PC – the values of squared cosines. Cosines refer to the correlation coefficients between the gene variables and PCs. The squared cosine reflects the representation quality of a gene variable on a PCA axis. (**F**) The PCA analysis on the standardized protein expressions of *EIF4G1, EIF4A1, EIF4E, EIF4EBP1, PABPC1, MKNK1, MKNK2* and *MYC* in 109 lung adenocarcinomas and 102 paired normal lung tissues from CPTAC.

**Fig. S8.**
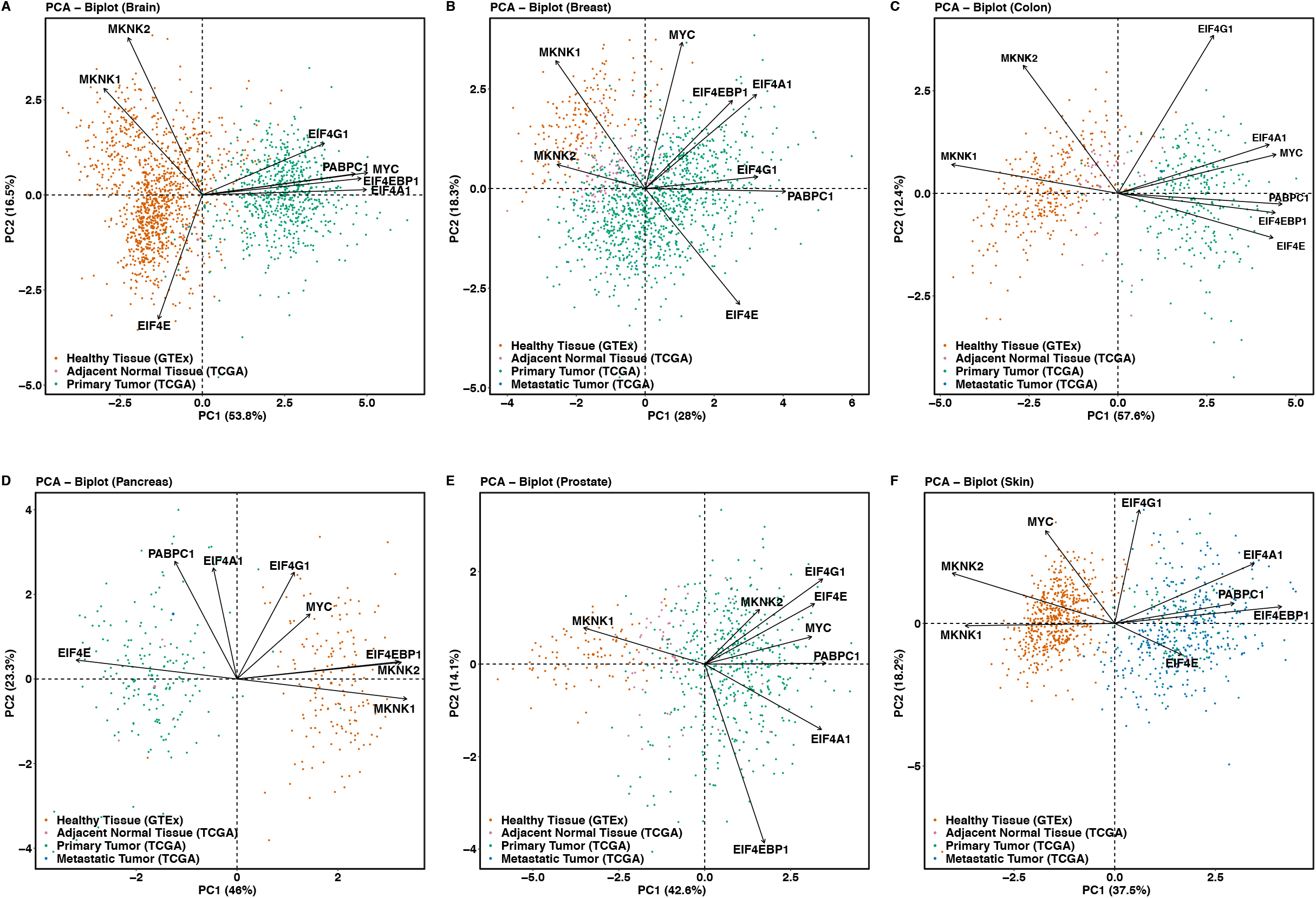
Malignant tumors are distinguished from healthy tissues by expressions of *EIF4F* subunits. (**A** to **F**) The PCA analysis using the standardized RNA expressions of *EIF4G1, EIF4A1, EIF4E, EIF4EBP1*, *PABPC1*, *MKNK1*, *MKNK2* and *MYC* from individual cancer types and the matched healthy tissues. The PCA biplot shows the first two PCs along which the samples vary the most. Axis title shows the percentage of variances (squared loadings) explained by the PC1 or the PC2. Arrows show the influence of each gene variable on the PCs. Axis values show the PCA scores of individual samples. The sample types were not used to construct PCs and they were labeled in color afterwards on the PCA plot. (**A**) The PCA analysis on 662 primary brain tumor samples (509 lower grade gliomas and 158 glioblastoma multiformes) and 5 adjacent normal brain tissues from TCGA, and 1,136 healthy brain tissues (including cerebellum, caudate, cortex, nucleus accumbens, cortex, hippocampus and hypothalamus) from GTEx. (**B**) The PCA analysis on 1,092 primary breast tumors, 7 metastatic breast tumors and 113 adjacent normal breast tissues from TCGA breast invasive carcinoma group, and 179 healthy breast mammary tissues from GTEx. (**C**) The PCA analysis on 288 primary colon tumors, 1 metastatic colon tumor and 41 adjacent normal colon tissues from TCGA colon adenocarcinoma group, and 304 healthy colon tissues (including transverse and sigmoid colon tissues) from GTEx. (**D**) The PCA analysis on 178 primary pancreatic tumors, 1 metastatic pancreatic tumor and 4 adjacent normal pancreas tissues from TCGA pancreas adenocarcinoma group, and 165 healthy pancreas tissues from GTEx. (**E**) The PCA analysis on 495 primary prostate tumors, 1 metastatic prostate tumor and 52 adjacent normal prostate tissues from TCGA prostate adenocarcinoma group, and 100 healthy prostate tissues from GTEx. (**F**) The PCA analysis on 102 primary skin tumors, 366 metastatic skin tumor and 1 adjacent normal skin tissue from TCGA skin cutaneous melanoma group, and 556 healthy skin tissues (including sun exposed and not sun exposed skin tissues) from GTEx.

**Fig. S9.**
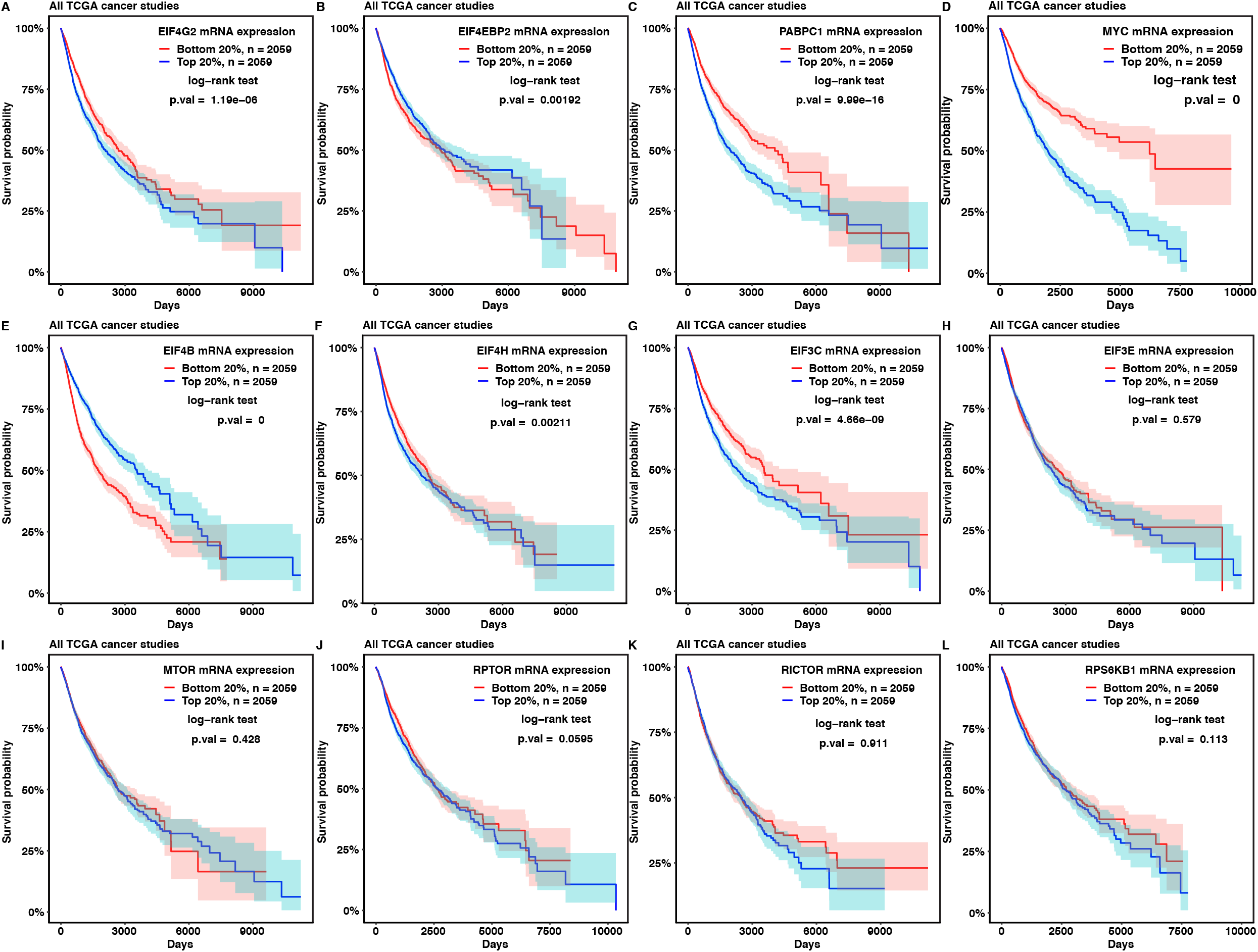
Survival analyses of *EIF4F* expression in patients from all cancer types. (**A** to **F**) The Kaplan-Meier plots compare the survival probabilities of TCGA cancer patients according to the mRNA expressions of *EIF4G2* (**A**), *EIF4EBP2* (**B**), *PABPC1* (**C**), *MYC* (**D**), *EIF4B* (**E**), *EIF4H* (**F**), *EIF3C* (**G**), *EIF3E* (**H**), *MTOR* (**I**), *RPTOR* (**J**), *RICTOR* (**K**), *RPS6KB1* (**L**) in their tumors. We ranked all 10,295 TCGA cancer patients based on the individual *EIF4F* gene expression from their tumor biopsies and selected two groups of patients with the top 20% or the bottom 20% of gene expression. The survival probability is the probability of an individual survives from the time origin (e.g. diagnosis of cancer) to a specified time. Differences in survival probabilities between two patient groups were assessed by the logrank test. The shaded areas around the associated survival curves show the lower and upper bounds of the 95% confidence interval.

**Fig. S10.**
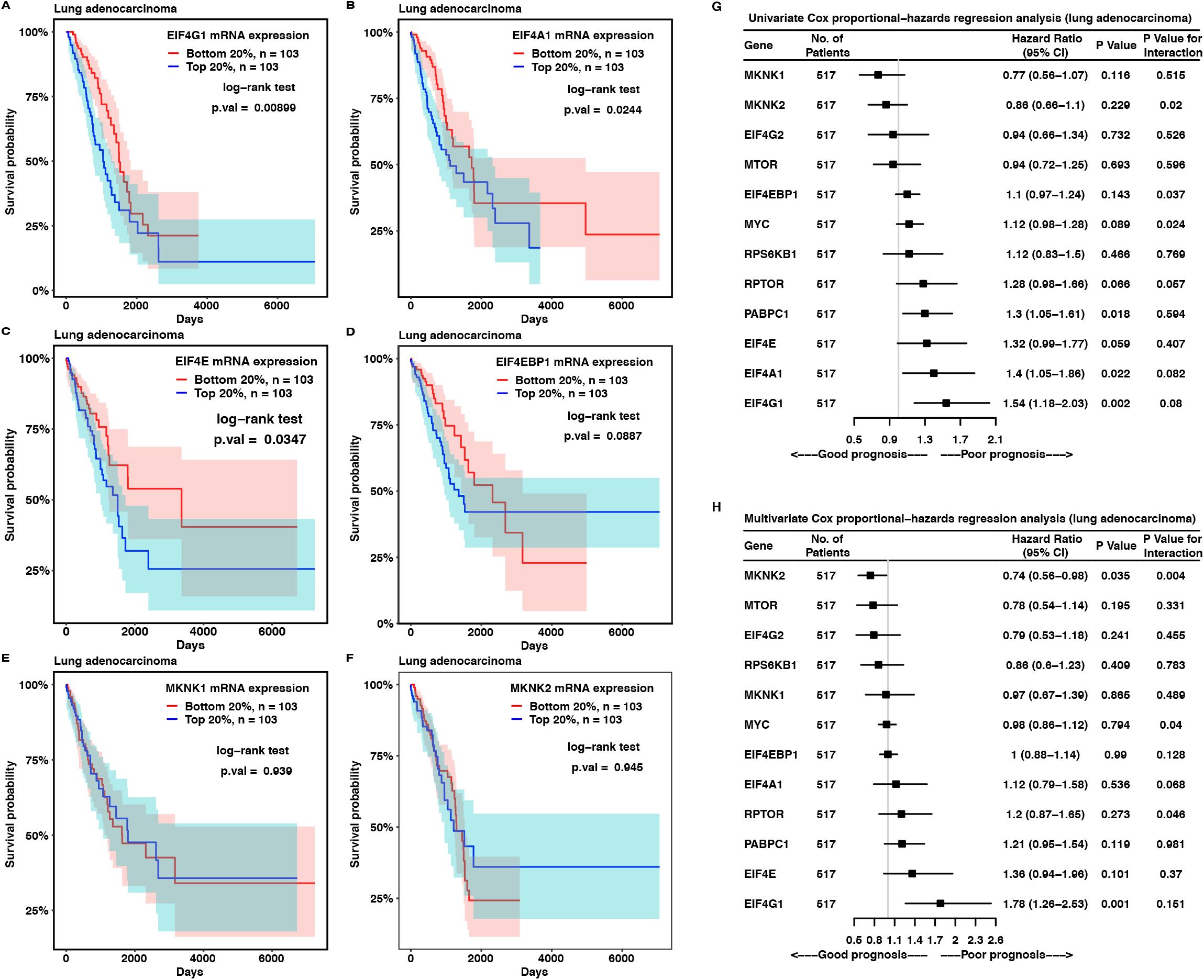
Survival analyses of *EIF4F* expression in lung adenocarcinoma patients. (**A** to **F**) The Kaplan-Meier plots compare the survival probabilities of lung adenocarcinoma patients according to the mRNA expressions of *EIF4G1* (**A**), *EIF4A1* (**B**), *EIF4E* (**C**), *EIF4EBP1* (**D**), *MKNK1* (**E**) and *MKNK2* (**F**). We ranked 517 TCGA lung adenocarcinoma patients based on the individual *EIF4F* gene expression from their tumor biopsies and selected two groups of patients with the top 20% or the bottom 20% of gene expression. Differences in survival probabilities between two patient groups were assessed by the log-rank test. The shaded areas around the associated survival curves show the lower and upper bounds of the 95% confidence interval. (**G** and **H**) The univariate (**G**) and multivariate (**H**) Cox proportional-hazards regression models were examined for *EIF4F* expression in 517 lung adenocarcinoma patients from TCGA. P value indicates the statistical significance of association between gene expression and survival (i.e. a significant fit in Cox-PH model). Assumption of Cox-PH model was examined by Schoenfeld residues to test constant hazard ratio over time. A statistically significant P value for interaction indicates a violation on proportional-hazards model. CI, confidence interval.

**Fig. S11.**
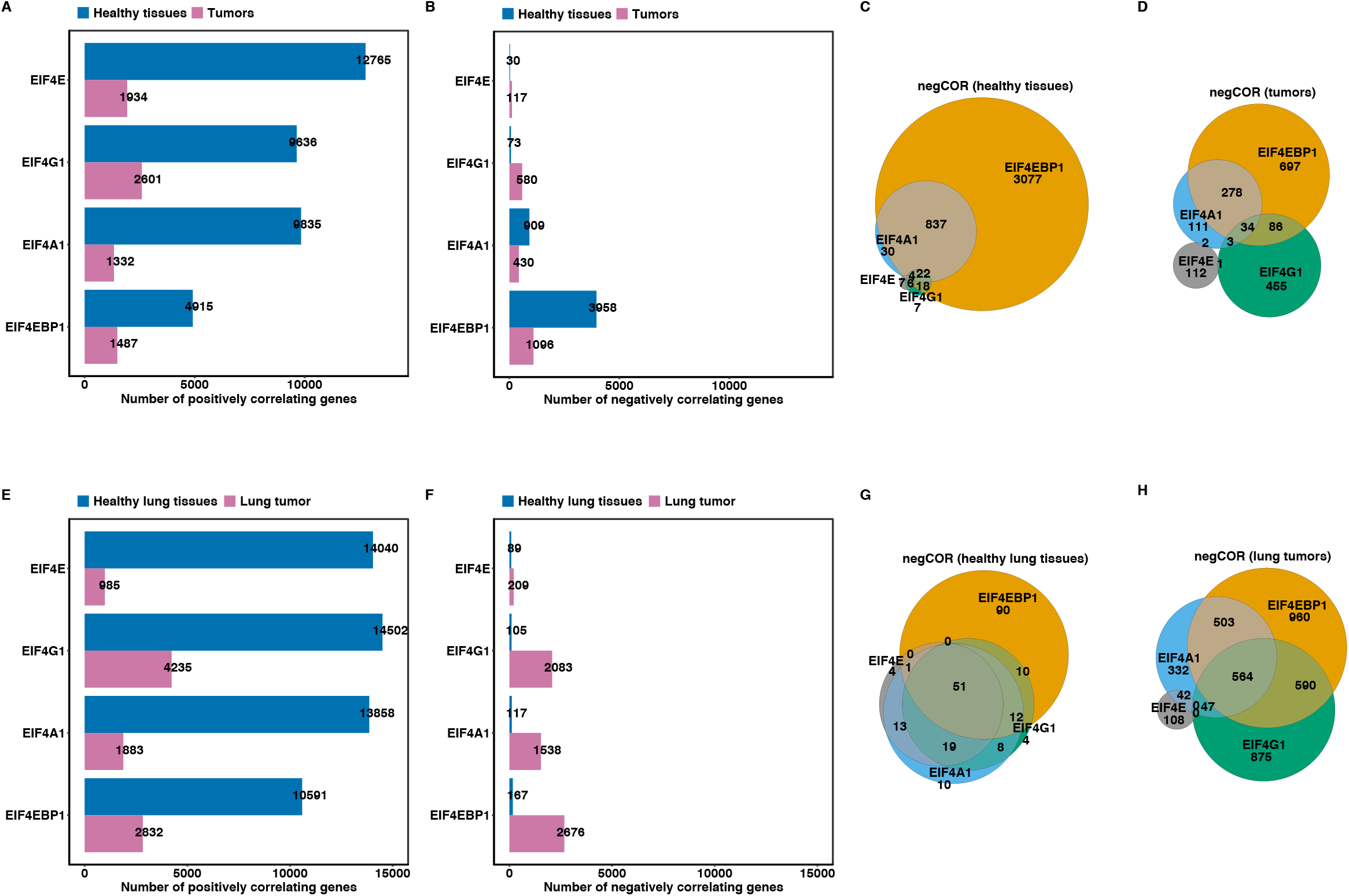
Tumors dysregulate genes that correlate with *EIF4G1* and *EIF4E* expressions. (**A** and **B**) Pearson’s correlation coefficients between *EIF4F (EIF4E, EIF4G1* or *EIF4A1*) and each of 58,582 genes were calculated separately from 10,323 TCGA tumor samples, or 7,414 GTEx healthy tissues, using the Toil Recomputed RNA-Seq datasets. The genes that have significant positive correlations (posCORs, r > 0.3) or negative correlations (negCORs, r < −0.3) were selected for analysis. The bar plots show the numbers of posCORs (**A**) and negCORs (**B**) identified for each *EIF4F* gene in tumors or in healthy tissues. (**C** and **D**) The Venn diagrams show the overlapping negCORs among *EIF4F* in healthy tissue samples (**C**), or in tumors (**D**). (**E** and **F**) The Pearson’s correlation coefficients between *EIF4F* (*EIF4E*, *EIF4G1*, *EIF4A1* or *EIF4EBP1*) and other 58,582 genes were calculated separately in 1,122 lung tumors of LUSC and LUAD from TCGA, or in 288 healthy lung tissues from GTEx, using the Toil recomputed RNA-Seq datasets. The bar plots show the numbers of posCORs (**E**) and negCORs (**F**) identified for each *EIF4F* gene from tumors and healthy tissues. (**G** and **H**) The Venn diagrams show the overlapping negCORs among *EIF4F* identified in healthy lungs (**G**), or in lung tumors (**H**).

**Fig. S12.**
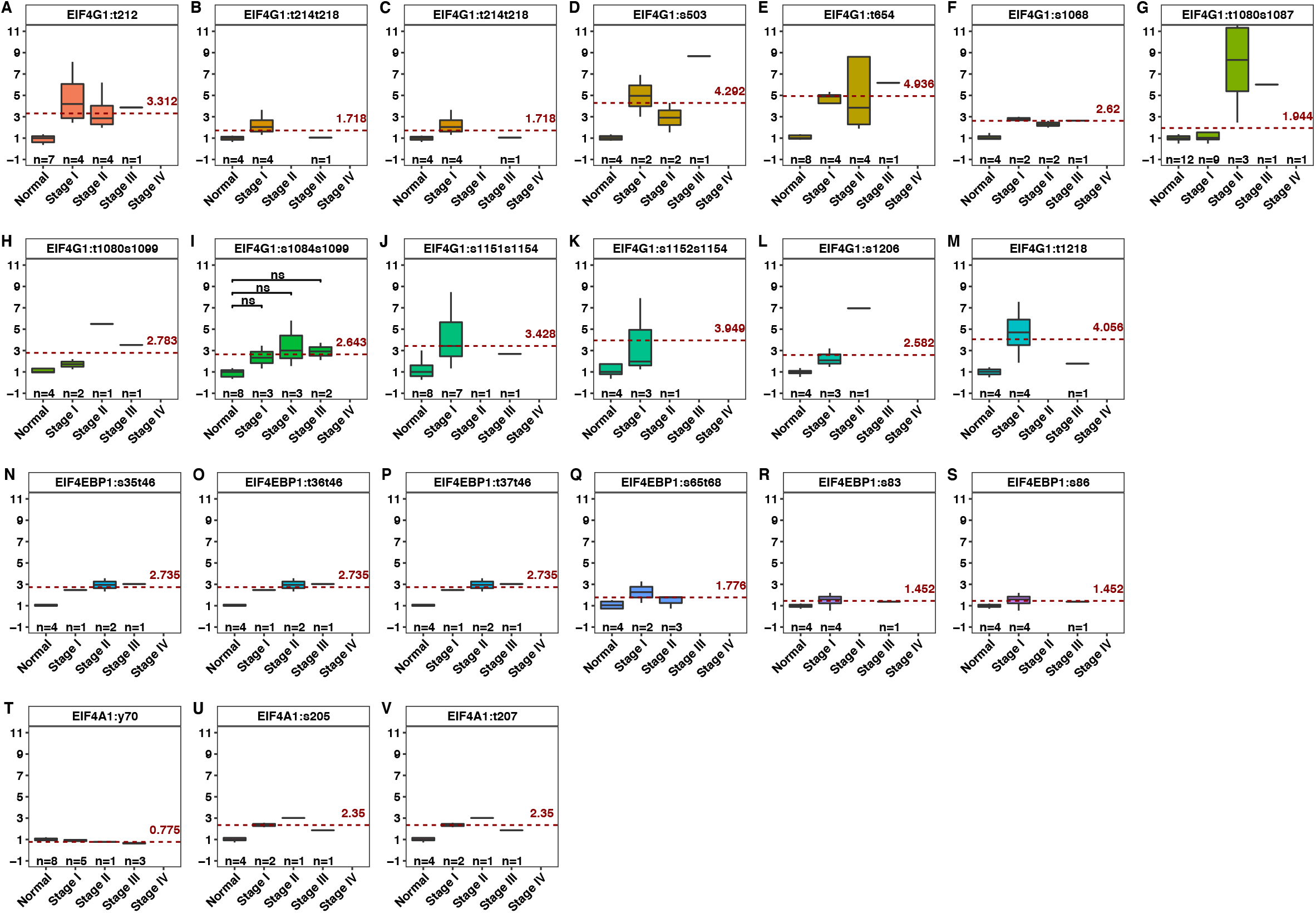
Phosphorylation of eIF4F subunits in lung adenocarcinoma. (**A** to **M**) The relative abundances of detected phosphor-peptides from eIF4G1 protein in normal lung tissues and different stages of lung adenocarcinomas. Y-axis indicates the relative abundance of a phosphor-peptide normalized to the median value of its abundance in normal lung tissues. s stands for serine residue and t stands for threonine residue. The red dash lines mark the average abundances of indicated phosphor-peptides in all lung adenocarcinomas, compared to those in normal lung tissues. The two-tailed Student’s t tests were performed. ns, not significant; *P ≤ 0.05; **P ≤ 0.01; ***P ≤ 0.001; ****P ≤ 0.0001 (**N** to **S**) The relative abundances of detected phosphor-peptides from 4E-BP1 protein from normal lung tissues and different stages of lung adenocarcinomas. (**T** to **V**) The relative abundances of detected phosphor-peptides from eIF4A1 protein from normal lung tissues and different stages of lung adenocarcinomas.

## METHODS

### Copy number variation analysis

Gene-level copy number data for 33 TCGA cancer types were obtained from the UCSC Xena data hub (Goldman et al., 2019) (https://tcga.xenahubs.net and https://pancanatlas.xenahubs.net). Genomic copy number variants from biopsies were measured by Affymetrix Genome-Wide Human SNP 6.0 arrays at the TCGA genome characterization center. The array data were normalized and estimated for raw copy number of genomic segments through the copy number variation (CNV) pipeline at NCI’s Genomic Data Commons (GDC). Gene-level CNVs were estimated from the copy numbers of genomic segments using the Genomic Identification of Significant Targets in Cancer 2.0 (GISTIC2) method (Mermel et al., 2011). GISTIC2 also grouped the estimated CNV values as homozygous deletion, heterozygous deletion, diploid, low-level gene copy gain, or gene copy amplification. We downloaded the CNV and annotation datasets from the UCSC Xena data hub.

For the correlation matrix of CNV among EIF4F genes, we used the TCGA pan-cancer gene-level copy number variation dataset that combined GISTIC2 analyzed data from all TCGA cohorts, with the Xena dataset ID: TCGA.PANCAN.sampleMap/Gistic2_CopyNumber_Gistic2_all_data_by_genes.

We used rcorr() function from the R package “Hmisc” to compute the correlation coefficients by Pearson correlation method and the p-value of the correlation.

To construct percentage stacked bar plots for illustrating CNV statuses of EIF4F subunits across TCGA cancer types, we used the TCGA pan-cancer gene-level copy number variation threshold dataset that combined GISTIC2-thresholded data from all TCGA cohorts, with the Xena dataset ID: TCGA.PANCAN.sampleMap/Gistic2_CopyNumber_Gistic2_all_thresholded.by_genes

To assess the correlation between CNV status and gene expression, we used the batch normalized RNA-Seq data published by The Cancer Genome Atlas (TCGA) Pan-Cancer analysis project (Schaub et al., 2018), with Xena dataset ID: EB++AdjustPANCAN_IlluminaHiSeq_RNASeqV2.geneExp.xena.

To analyze CNV ratios between tumor and normal tissues across TCGA cancer types, we used the copy ratio data between tumor and normal tissues that were generated from the Affymetrix SNP6.0 array data by Tangent Normalization method (Tabak et al., 2019), with the Xena dataset ID: broad.mit.edu_PANCAN_Genome_Wide_SNP_6_whitelisted.gene.xena.

We acquired the clinically relevant phenotype information for each TCGA sample, including sample type and primary disease annotation datasets from all individual TCGA cohorts, with the Xena dataset ID: TCGA_phenotype_denseDataOnlyDownload.tsv.

### RNA-Seq data for gene expression analysis

The original underlying mRNA sequencing of tumor samples from TCGA, and healthy samples from GTEx, were obtained for those organizations. All RNA-Seq experiments used the poly(A) enrichment method for RNA preparation. In order to treat data from those two sources in a consistent way and minimize computational batch effects on read alignment and quantification, we used the reprocessed RNA-Seq read count data for both sources, available from the UC Santa Cruz computational genomics Lab, computed with the Toil-based RNA-Seq bioinformatic pipeline (Vivian et al., 2017). In the Toil-based pipeline, paired-end reads of RNA-seq samples from the TCGA and GTEx projects were processed by STAR to align sequence reads to the GRCh38/hg38 human reference genome and generate read coverage, and by RSEM to quantify number of RNA-Seq reads that aligned to a transcript. Transcript-level expression was estimated as transcripts per million (TPM), which divided the mapped RNA-Seq reads by the length of transcript to give transcript-level expression independent of transcript length. Genelevel expression data were calculated by summing up all transcript-level TPM for each gene. Gene-level expression data were quantile normalized to facilitate comparison across samples and experiments, and then log_2_(x+1) transformed to remove extreme values. Gene-level expression datasets were presented in terms of the counting unit: log_2_(normalized_TPM+1). We used the sample annotation dataset for clinically relevant phenotype information of each sample, including tumor primary site, tumor subtypes, and primary tissues. We obtained the gene expression and sample annotation datasets from UCSC Xena data hub (https://toil.xenahubs.net), with the Xena dataset IDs: TcgaTargetGtex_RSEM_Hugo_norm_count, and TcgaTargetGTEX_phenotype.txt.

To compute the sum expression or expression ratios, we back-transformed counting units to obtain the normalized TPM values for each gene.

### Principal component analysis

PCA analysis on RNA-Seq was performed on the combined gene expression data from TCGA and GTEx by variances in gene expressions (log_2_(normalized_TPM+1)). The gene expression data were scaled to even out the variances between principal components (PCs) prior to performing the PCA transformation. We used the function PCA() implemented in the R package “FactoMineR” to draw PCA biplots. We used the function get_pca_var() to extract cos2 of gene variables, and corrplot() (from “corrplot” R package) to visualize representation of the gene variable on each PC.

Among 33 TCGA cancer types, there are eighteen study groups containing fewer than 10 matched adjacent normal tissues from cancer patients, which makes statistical comparison between tumor and normal samples difficult. When performing PCA analysis on all cancer types, we only included TCGA tumor samples (primary and metastatic tumors) and GTEX normal tissue samples from healthy individuals. When we performed PCA analysis on individual cancer types, we included tumor samples and two types of normal samples: TCGA’s solid adjacent normal tissues from cancer patients, and matched GTEx’s normal tissue from healthy individuals.

PCA analysis on proteomics was performed on the relative protein abundance data from CPTAC with a variance in log_2_(ratio). The proteomics data were scaled to even out the variances between PCs prior to performing the PCA transformation. Because the CPTAC proteomics dataset missed expression values for some proteins, we used the regularized iterative PCA algorithm to impute the missing expression data. We used the function impute.PCA() from the R package “missMDA” (two dimensions were chosen) to impute the proteomics dataset, and then performed PCA on output dataset using the PCA() function from “FactoMineR” package.

### Survival analysis

Tumors from 11,160 patients of 33 different cancer types were collected at TCGA. Those tumors were originally diagnosed from 1978 to 2013 and mostly primary tumors (except skin cutaneous melanoma). For primary tumors, the biopsies were removed from the patient at or close to the time of diagnosis. Overall Survival (OS) data were selected as the clinical endpoints for pan-cancer survival analysis. OS was defined as the period from the date of diagnosis until the date of death from any cause, with the date of diagnosis chosen as time zero.

We obtained the relevant datasets from UCSC Xena data hub (https://pancanatlas.xenahubs.net). We used the curated clinical data generated by the TCGA Pan-Cancer Clinical Data Resource (Liu et al., 2018), with the Xena dataset ID: Survival_SupplementalTable_S1_20171025_xena_sp. We used the batch normalized RNA-Seq data from TCGA Pan-Cancer atlas (Schaub et al., 2018), with the Xena dataset ID: EB++AdjustPANCAN_IlluminaHiSeq_RNASeqV2.geneExp.xena. We excluded all samples that were annotated as “solid tissue normal” from survival analysis, using the sample type annotation dataset with the Xena dataset ID: TCGA_phenotype_denseDataOnlyDownload.tsv. The unit for gene expression was log_2_(norm_value+1) in the expression dataset, which was used for the survival analyses. We examined the data distribution of twelve *EIF4F* relevant genes (log_2_(norm_value+1)) by density plots. Although RNA-Seq data do not form normal distribution, all twelve genes displayed bell-shaped curves without obviously extreme values across 10,295 tumor samples.

For Kaplan-Meier analysis on gene expression, two subgroups (top and bottom 20% groups) out of cancer patients were selected, according to *EIF4F* expression within their tumors. We created the survival plots for two subgroups using survfit() function and compared the difference of survival curves by the logrank test using survdiff() function from the R package “survival”.

The Cox proportional hazards (PH) regression model was used to develop a predictive model of overall survival, based on the gene expression values and clinical outcomes from TCGA cancer patients. We used the coxph() function to compute Hazard Ratio (HR), 95% confidence interval, and statistical significance for gene expression in relation to overall survival (P value). The proportional hazards PH assumption was assessed by correlating the Schoenfeld residuals and time to test for dependence, using the cox.zph() function from the “survival” package. For each gene, the statistical significance for relationship between Schoenfeld residuals and time is shown as the P value for interaction. A significant P value for interaction (< 0.05) indicates the PH assumption is violated.

### Correlation analysis and Venn diagram

The linear dependence between each of *EIF4F* genes and all other genes identified by RNA-Seq (58,582 genes in total) were measured by Pearson’s correlation method in 10,323 TCGA tumor samples, or in 7,414 GTEx healthy tissues. For consistency, we used the reprocessed RNA-Seq data for the TCGA and GTEx datasets with the Toil-based pipeline. Gene expression data for correlation analysis were in the count unit as log_2_(normalized_TPM+1). We used the cor.test function in R to measure the association between paired genes, with the Pearson’s correlation coefficient (r) and statistical significance of the correlation (p-value). We selected the genes with r values greater than 0.3 (p-value ≤ 0.05) as the significant positive correlation genes (posCORs), and the genes with r values less than −0.3 (p-value ≤ 0.05) as the significant negative correlation genes (negCORs). Venn diagram was used to illustrates levels of overlap between correlating genes. We used VennCounts() function from the R package “limma” to count the overlapping genes between gene groups, then used the counts to draw the proportional Venn diagrams with euler function from the R package “venneuler”.

### Heatmap, clustering and pathway enrichment analysis

The heatmap was created to visualize the Pearson’s correlation coefficients of posCORs and negCORs for *EIF4E, EIF4G1, EIF4A1* and *EIF4EBP1* from tumors or normal samples. Using Heatmap() function of the R package “ComplexHeatmap”, we drew heatmaps using rows to represent posCORs and negCORs. The heatmap rows were ordered and partitioned into three non-overlapping subgroups by K-means clustering method (k = 3). The heatmap columns represent *EIF4F* genes and sample types. The columns were ordered by the hierarchical clustering method. The gene list within K-means cluster from heatmap was retrieved for pathway enrichment analysis.

The pathway-based analysis was performed on the clustered gene lists with the enrichPathway() function from the R package “ReactomePA”, which used Reactome as a source of pathway data. Statistical analysis and visualization of enriched biological pathways were performed with the compareCluster() function from R package “clusterProfiler”, by over-representation analysis (ORA) method. The statistical significance (p-value) of the overlap between genes from a given pathway and the clustered gene list was determined by the hypergeometric distribution test. The p-values were adjusted for multiple comparison by the Hochberg’s and Hommel’s method, using p.adjust function (from the R package “stats”) with method = “BH”. The enriched pathways were ordered according to their adjusted p-values. The ratios between the number of genes associated with a given pathway and the total number of genes in the clustered list were calculated as gene ratios.

### Proteomics and phosphor-proteomics data for phosphorylation analysis

Proteomics and phosphor-proteomics data of 109 lung adenocarcinomas (LUAD) and 102 paired normal adjacent tissue samples were generated by the Clinical Proteomic Tumor Analysis Consortium (CPTAC). An isobaric peptide labeling approach (iTRAQ) was employed to quantify protein and phosphor-peptides across samples. For proteomics, protein extraction from biopsies were labeled with 10-plex tandem mass tags (TMT) reagents and analyzed by liquid chromatography with tandem mass spectrometry LC-MS/MS (Mertins et al., 2016). For phosphor-proteomics, phosphor-peptides were enriched by immobilized metal-ion affinity chromatography and then analyzed by LC-MS/MS. The phosphorylated amino-acids within peptides were mapped to the corresponding proteins’ sites. To facilitate quantitative comparison between all samples across experiments, a pooled reference sample from all samples was included in each 10-plex experiment. All data processed steps including peptide sequence alignment to proteins and peptide quantitation were performed with the common data analysis pipeline (CDAP) at CPTAC.

Expression data were obtained from CPTAC_LUAD_Proteome_CDAP_Protein_Report.r1 and CPTAC_LUAD_Phosphoproteome_CDAP_Protein_Report.r1 at the CPTAC website. The sample annotation data were obtained from CPTAC_LUAD_metadata.

### Quantification and Statistical Analysis

Statistics were performed on R (version 3.6). The p-value for Pearson’s correlation was calculated using a t-distribution with n – 2 degrees of freedom. Unpaired Student’s t tests or one-way analysis of variance (ANOVA) were used to compare the expression data. The p-value for Kaplan-Meier analysis was determined by the log-rank test. The p-value for Cox regression analysis was determined by the likelihood-ratio test. The p-value for the pathway enrichment analysis was determined by the hypergeometric distribution test and further adjusted for multiple comparison by the Hochberg and Hommel method. p-values < 0.05 were considered statistically significant.

